# Genotypic and phenotypic diversity of *Maudiozyma humilis*: the multiple evolutionary trajectories of a domesticated yeast

**DOI:** 10.64898/2026.02.16.706159

**Authors:** Manon Lebleux, Jess Rouil, Diego Segond, Thérèse Marlin, Kate Howell, Pamela Bechara, Thibault Nidelet, Lucie Arnould, Delphine Sicard, Hugo Devillers

## Abstract

*Maudiozyma humilis* is the second most frequently encountered yeast species in sourdough bread. Despite its ecological and food relevance, little is known about its evolutionary trajectories and phenotype traits of interest. Here we investigated the genomic and phenotypic diversity of a world-wide collection of 55 *M. humilis* strains, including 52 from sourdough, by combining genomic analysis, flow cytometry and high-throughput phenotyping of fermentation kinetics and fitness.

Population genomic analysis revealed six genetically distinct clades, three diploid and three triploid, with no geographical or substrate-specific structuring. Phylogenetic, loss of heterozygosity (LOH) distribution and allele specific analysis indicated that triploid strains originated from both recent and more ancient hybridization events involving multiple diploid lineages. The absence of the *HO* gene, and mating-type silent cassettes, revealed that *M. humilis* is not able to carry mating-type switching. In addition, high linkage disequilibrium (LD) and variable LOH accumulation were observed consistent with a predominantly clonal reproduction. Last, an exceptionally high level of heterozygosity was detected, suggesting that occasional hybridization is the major driver of genetic diversity.

Phenotypic characterization in a synthetic sourdough medium revealed variation in fermentation kinetics and fitness statistically associated with the genetic clades. Interestingly, genetic distance between clades and strains better explain the phenotypic variation than difference in ploidy.

Altogether, our findings highlight the complex evolutionary history of *M. humilis*, shaped by hybridization and ploidy variation and reveal that historical contingency, more than ploidy, shapes the phenotypic landscape of this species. Beyond providing the first analysis of *M. humilis* evolution, our result challenges the hypothesis that increases in ploidy are necessarily beneficial in domesticated species.

## Introduction

The development of fermented foods has resulted from a multicultural process, ultimately yielding the diversity of products observed today. During this long process, humans have shaped the evolution of a wide diversity of yeast species, which are responsible for many fermentation processes. However, beyond the well-known *Saccharomyces* species, relatively little is known about the evolution of other yeast species commonly involved in food fermentation. Yet, many yeast species have been described in fermented products, particularly in naturally fermented artisanal products that do not involve the addition of strains artificially selected by humans for industrial purposes (Hittinger et al. 2018). This is the case of yeast species found in sourdough bread-making where more than 80 yeast species belonging to more than 22 genus have been detected (De Vuyst et al. 2016; Carbonetto et al. 2018; Arora et al. 2021; Calvert et al. 2021; von Gastrow, Gianotti, et al. 2023).

Sourdough is a mixture of flour and water inhabited by a community of bacteria and yeast species, typically including one to three dominant lactic acid bacteria species and one dominant yeast species (Arora et al. 2021; Calvert et al. 2021; von Gastrow et al. 2023). Although *Saccharomyces cerevisiae* is the most frequently encountered dominant yeast species in sourdough, several other yeast species belonging to different genera within the subphylum Saccharomycotina are also regularly found as dominant, including species of the genera *Maudiozyma*, *Monosporozyma*, *Pichia, Torulaspora*, and *Wickerhamomyces* (De Vuyst et al. 2016; Carbonetto et al. 2018; Arora et al. 2021; Calvert et al. 2021; von Gastrow et al. 2023). Among these genera, the genus *Maudiozyma* (formerly *Kazachstania*), a closely related genus of *Saccharomyces*, is by far the most represented genus in sourdough. Eight *Maudiozyma* species have been described in bread sourdough, namely *Maudiozyma humilis*, *Maudiozyma bulderi*, *Maudiozyma barnettii*, *Maudiozyma exigua, Maudiozyma pseudohumilis, Maudiozyma saulgeensis, Maudiozyma australis* and *Maudiozyma bozae* (Jacques et al. 2016; Sarilar et al. 2017; Gouliamova and Dimitrov 2020; Landis et al. 2021; Bazalová et al. 2022; Devillers et al. 2022; Wittwer et al. 2022; Michel et al. 2023; Liu et al. 2024). Among these species, *M. humilis* is the second most frequently isolated yeast species after *S. cerevisiae* in sourdough (Carbonetto et al. 2018; Arora et al. 2021; Calvert et al. 2021; von Gastrow et al. 2023).

*M. humilis* has been found in home-made and bakery sourdoughs around the world (Urien et al. 2019; Boyaci-Gunduz and Erten 2020; Chiva et al. 2020; Comasio et al. 2020; Wang et al. 2020; Landis et al. 2021; von Gastrow, Gianotti, et al. 2023; Michel et al. 2023; Bai et al. 2024; Dessalegn and Andualem 2024; Sanmartín et al. 2024). In France, its occurrence is linked to bread-making practices, being more frequent in sourdoughs produced with artisanal bakers than in those made by farmer-bakers, who bake bread less frequently and in smaller quantities using flour most often made on farm from stone-milled grains of wheat landraces (Michel et al. 2023) To date, *M. humilis* has predominantly been isolated from sourdough environments, with occasional reports in other fermented products, such as coffee (Qin et al. 2024; Suo et al. 2025), cocoa (Papalexandratou et al. 2019; Velásquez-Reyes et al. 2025), peach waste (Gonçalves et al. 2024), pepper (Li et al. 2024), sugarcane bagasse (Peng et al. 2021), baijiu (You et al. 2021; Dong et al. 2025; Lin et al. 2025), fish (Punyauppa-path et al. 2022), banana (Mure et al. 2023), agave (García-Ortega et al. 2022; Colón-González et al. 2025), chilli sauce (Feixia et al. 2024), pickled chayote (Shang et al. 2022), miang (fermented tea leaf product) (Phovisay et al. 2024), msalais (fermented boiled grape) (Zhu et al. 2023), tarhana (Ozel et al. 2025), Sichuan *shai* vinegar (Zhu et al. 2026). *M. humilis* has also been found in non-food fermentation environments, such as maize and sorghum silages (Lai et al. 2025; Tahir et al. 2025), bio-hydrogen production using dark fermentation (Detman et al. 2018) and human fecal microbiome (Vonaesch et al. 2023). This species has rarely been detected in other environments than fermented products. One study reported the presence of *M. humilis* in marine environment in China (Xue et al. 2024) and another on tree bark and rhizosphere in Ethiopia (Dabassa Koricha et al. 2019).

The genomes of nine *M. humilis* strains are sequenced, including 8 available on NCBI. The type strain has been sequenced twice (Opulente et al. 2024), it was isolated from Bantu beer in South Africa. Four strains originated from agave fermentation in Mexico (García-Ortega et al. 2022; Gallegos-Casillas et al. 2024). The three remaining strains were isolated from fermented banana in Japan (Mure et al. 2023), sourdough in the United States (Opulente et al. 2024), and bioreactors in the sugar industry in Poland (Detman et al. 2018; Mielecki et al. 2024), respectively. Assemblies ranged from 13.25 to 19.11 Mb with an average GC content of 49%. However, genomic data on *M. humilis* remains limited. Ploidy level has been reported only for the diploid strain, MAW1 (Mielecki et al. 2024) and genome annotation is available only for strain KH-74 (Mure et al. 2023).

Regarding phenotypic aspects, only a few studies have been made on *M. humilis* involving a limited number of strains. Convergent results have highlighted the tolerance of this species to various stresses, such as 9% ethanol and up to 10% w/v of NaCl and its ability to metabolize fructose, glucose, galactose, trehalose and glycerol but not raffinose, xylose, lactose. However, *M. humilis* was not described as able to consume maltose, unlike *S. cerevisiae*. As for sucrose and mannose, results diverged on its ability to consume them. Moreover, its sensitivity to low pH levels may vary compared to *S. cerevisiae* in a strain-dependent manner even though it seemed to be more tolerant to acetic acid than *S. cerevisiae* (Brandt et al. 2004; Jacques et al. 2016; Carbonetto et al. 2020; Wittwer et al. 2022; Sánchez-Adriá et al. 2023; Mielecki et al. 2024; Yu et al. 2025).

The relative fermentation performance of *M. humilis* compared with *S. cerevisiae* varies across studies, sometimes described as inferior to *S. cerevisiae* (Xu et al. 2019; Carbonetto et al. 2020; Boudaoud et al. 2021), sometimes described as at least as good (Carbonetto et al. 2020; Sánchez-Adriá et al. 2023; Sanmartín et al. 2024), notably with regards of CO_2_ production. Compared to other yeast species, *M. humilis* was found to outperform *M. bulderi* or *Wickerhamomyces anomalus* (Xu et al., 2019, Michel et al., 2023). Globally these studies suggest an important phenotypic variability between strains of this species.

Therefore, *M. humilis* is an interesting model yeast species to gain further insight into the evolutionary trajectories of yeasts involved in fermented products. However, to fully understand the evolution and exploit this species, genomic and phenotypic diversity needs to be examined. Here, we explored the genomic and phenotypic diversity of a wide collection of 55 strains of *M. humilis* species, 52 originating from sourdough. Using whole genome sequencing, ploidy characterization by flow cytometry, high-throughput fermentation and fitness analysis, we investigate the evolutionary history of this sourdough yeast species and explore links between genomic and phenotypic landscapes.

## Results

### Complete genome sequences of 55 *M. humilis* strains

We sequenced the complete genomes of 55 *M. humilis* strains isolated from 18 different countries, representing five continents: Europe, Oceania, Asia, Africa and North America (Table S1). The acquired Illumina short-reads, whose coverage ranged between 92X and 357X (Table S2), have been deposited to the SRA database under the BioProject PRJEB89020.

### Polyloidy and chromosomal structural variations

Flow cytometry was used to assess the ploidy of the 55 *M. humilis* strains. A total of 28 strains were diploids, while the remaining 27 strains were triploids (Table S1). In order to identify large structural variations in the genomes of the 55 *M. humilis* strains, sequenced reads were mapped against a reference genome and coverages were investigated. This revealed large redundant regions in the reference genome used (Fig. S1), which have been masked in the subsequent analyses of read coverages.

Analysis of read coverage showed that most of the contigs had a uniform read distribution, although several of them had significant coverage variations, revealing both segmental duplications and segmental losses (Table S2). For example, in strain CBS 5658^T^, the beginning of contig 3 and the whole contig 15 had lost 50% of the representative reads, indicating that these regions were haploid (Fig. 1A). By contrast, contigs 11 and 14 showed a 50% increase in read coverage, suggesting they were present in three copies in the genome of this strain. Finally, the two-fold increase of read coverage at the end of contig 6 showed that this genomic fragment was present in four copies (Fig. 1A). Overall, the analysis of the 55 *M. humilis* strains revealed that most of the segmental variation were found at contig boundaries (vertical dashed grey lines in Fig. 1B). Five strains have segmental variations of complete contigs, namely LbY2, LbrY1, MTF 5052, CBS 2664 and CBS 5658^T^, suggesting possible aneuploidy. Last, some variations appeared to be shared by certain strains, such as about two-third of the way along the contig 3, at the beginning of contig 4 or the middle of contig 9 (Fig. 1B).

**Fig. 1:**
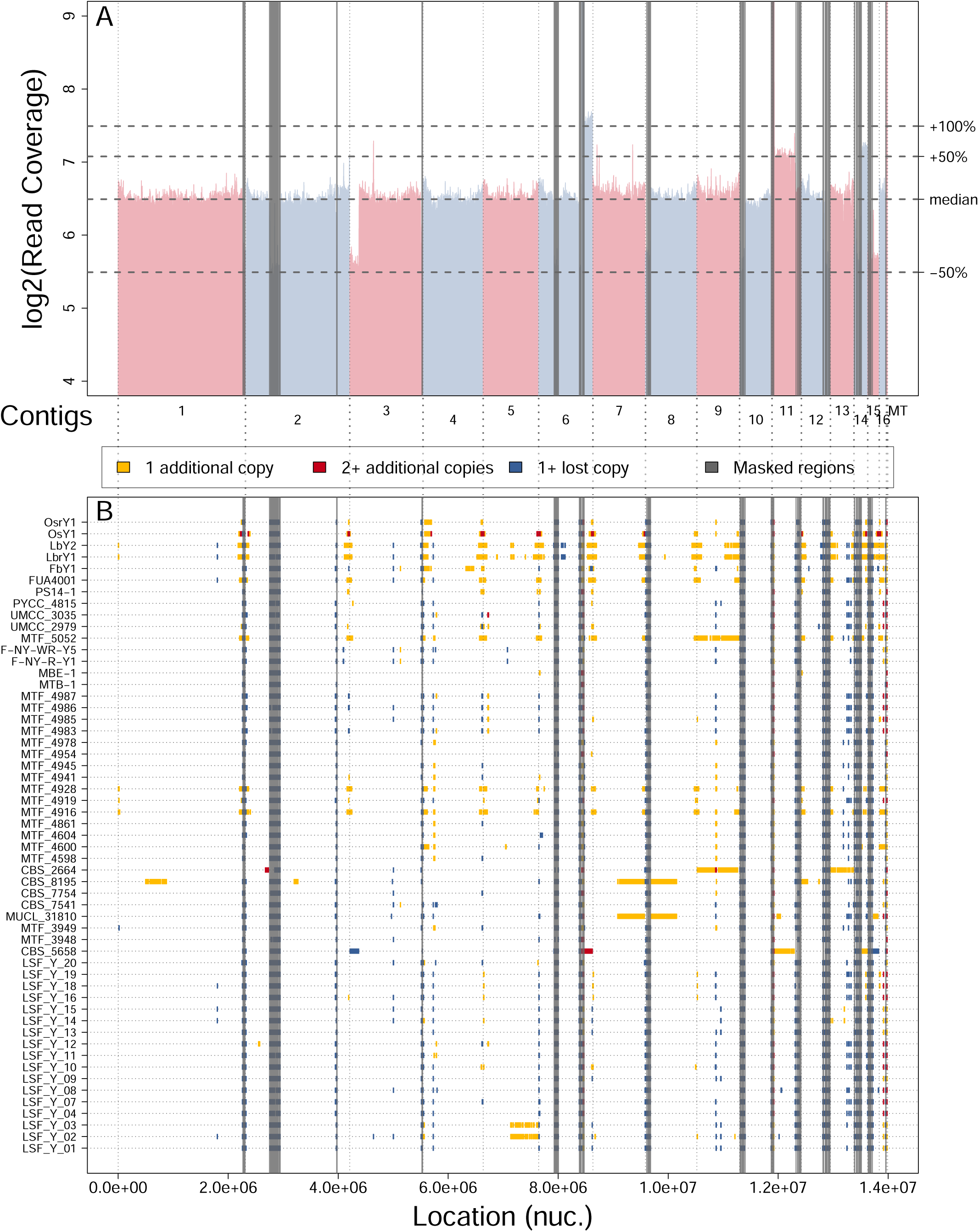
Identification of segmental variations in *M. humilis* genomes. (A) Read coverage of the diploid strain CBS 5658*^T^* along the reference genome (in Log2 scale). The +50% and +100%, which express an increase in coverage relative to the median value, represent an acquisition of one and two segmental copies, respectively. The -50% threshold corresponds to a loss on one segmental copy. (B) Summary of identified segmental variations of at least 10kb in all the 55 *M. humilis* strains based on the analysis of read coverage variation.

### All strains of *M. humilis* are highly heterozygous

A variant calling analysis was performed on the 55 strains of *M. humilis*, revealing a total of 1,565,767 distinct variant positions, including single nucleotide polymorphisms (SNPs) and insertions/deletions (INDELs). Biallelic SNPs represented a total of 1,189,761 sites among the 55 strains of *M. humilis*, with the distribution per strain ranging between 188,409 (strain CBS 5658^T^) and 542,037 (strain MTF 4604) (Fig. 2A and Table S3). Note that the strain CBS 5658^T^ had no homozygous SNPs because it is similar to the CLIB 1323^T^ used as reference genome in this study (Fig. 2A).

**Fig. 2:**
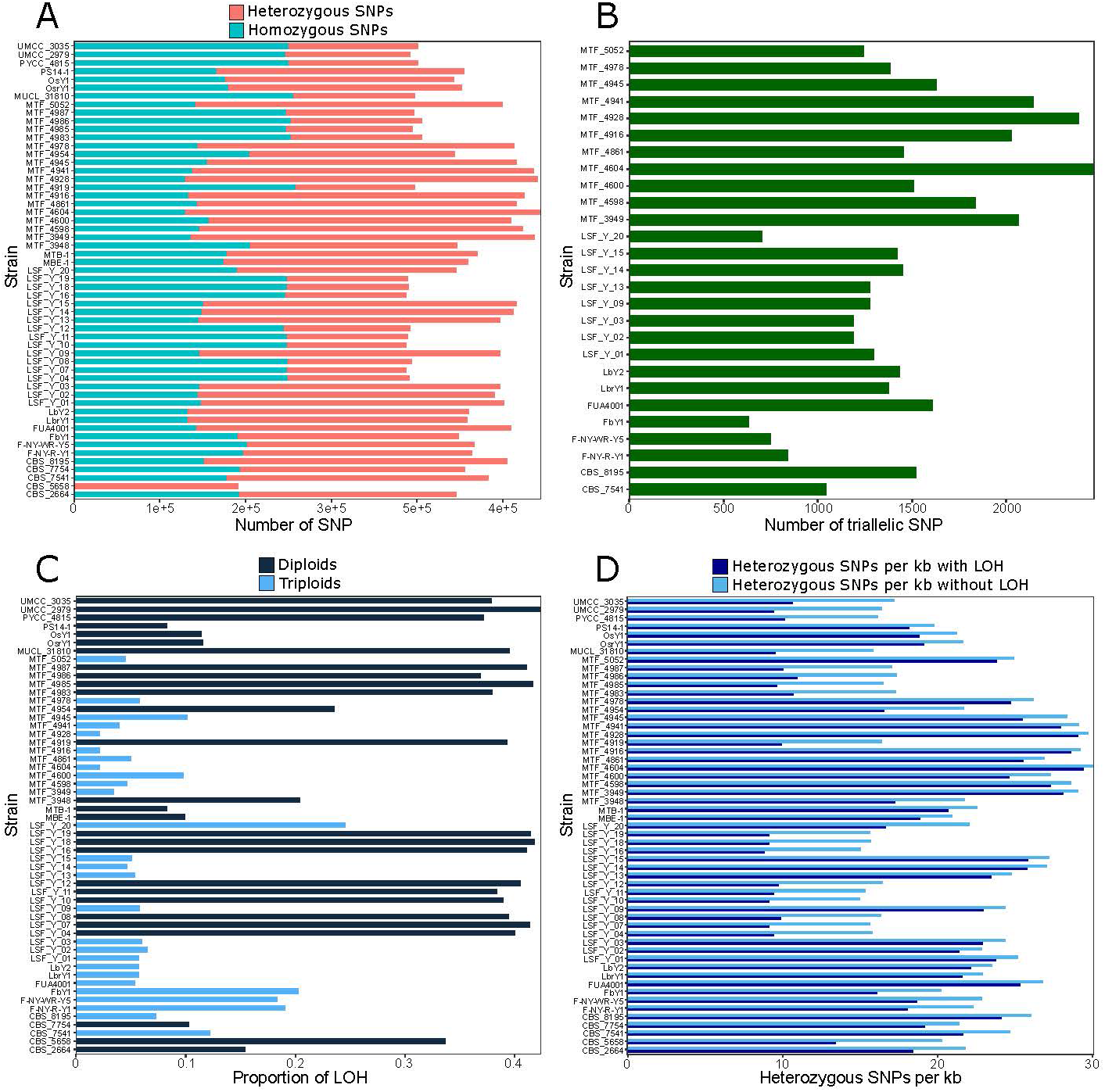
Distribution of homozygous and heterozygous SNPs and loss of heterozygosity (LOH) accumulation across the 55 *M. humilis* strains. (A) Number of homozygous SNPs (blue) and heterozygous SNPs (red) in all strains. (B) Number of heterozygous tri-allelic SNPs in triploid strains. (C) Proportion of the genome with LOH for all strains. (D): Number of heterozygous SNP per kb for all strains.

Among the selected biallelic SNPs, the number of heterozygous sites per strain ranged from 124,231 (strain LSF_Y_16) to 412,451 (strain MTF 4604) (Fig. 2A and Table S3). The proportion of heterozygous SNPs in diploid strains ranged between 35.1% and 63.7% while in triploid strains it varied from 56.9% to 72.2%. (Fig. 2A and Table S3). For the 27 triploid strains, triallelic heterozygous sites (*i.e.*, three distinct alleles at one site) were investigated. Between 469 (strain FbY1) and 1506 (strain MTF 4604) triallelic SNPs were identified (Fig. 2B), representing less than 0.2% of the total number of SNPs found on average in these strains (Table S3).

The variation of heterozygous sites in the 55 strains of *M. humilis* was further analysed by identifying regions of loss of heterozygosity (LOH) using the JLOH tool (Schiavinato et al. 2023). Between 15 and 336 LOH regions across the different genomes, covering from 302 kb to 5,933 kb, were identified depending on the strain (Table S4). This represented between 2.2% and 42.4% of the length of the reference genome (Table S4 and Fig. 2C). The size of individual LOH regions varied widely, ranging from the chosen detection threshold (3001 bases) to 857 kb depending on the strain (Table S4).

Heterozygosity rates were compared across strains, considering either the whole genome or the part of the genome without LOH (Fig. 2D and Table S3). When considering the whole genome length, the heterozygosity rate ranged between 8.9 SNPs/kb (strain LSF_Y_16) to 29.6 SNPs/kb (strain MTF 4604). When removing LOH, it reached values between 15.0 SNPs/kb (strain LSF_Y_10) and 30.2 SNPs/kb (strain MTF 4604).

### *M. humilis* population is structured into six clades

Population structure of the 55 strains of *M. humilis* was inferred using three distinct approaches, namely ADMIXTURE, DAPC and a phylogenomic reconstruction. To do so, the collection of biallelic SNPs was refined to remove any sites that were implied in a LOH region in at least one strain. Indeed, while a LOH region may arise from a single molecular event it can impact hundreds and even thousands of heterozygous sites leading to critical bias in population structure studies. A total of 72,561 biallelic SNPs were then retained.

The model-based clustering algorithm of ADMIXTURE tool (Alexander et al. 2009) was first used to infer the population structure based on the 72,561 retained SNPs. The cross-validation procedure suggested an optimal clustering at K = 6 clusters (Fig. S2A). Analysis of ADMIXTURE predictions for K = 6 and K = 7 revealed a robust genetic differentiation into 6 clades for 53 strains.

Two strains (CBS 5658^T^ and LSF_Y_20) were predicted with mixed ancestries when K = 6 and clustered in a distinct seventh clade when K = 7 (Fig. 3). These clades coped well with strain ploidy, clades 1, 3 and 6 contained only triploid strains while clades 2, 4 and 5 contained only diploid strains (see K = 6, Fig. 3). Assuming K = 5, a group of eleven triploid strains showed an admixture consisting of about two-thirds contribution from diploid clade 2 and one-third contribution from triploid clade 6 (Fig. 3). Interestingly, when considering K = 6 or K = 7, these eleven strains form a distinct cluster denoted clade 1. Among this latter, four strains (CBS 8195, FUA4001, LbrY1, and LbY2) exhibited minimal admixture, with less than 10% contribution from clade 2 (Fig. 3).

**Fig. 3:**
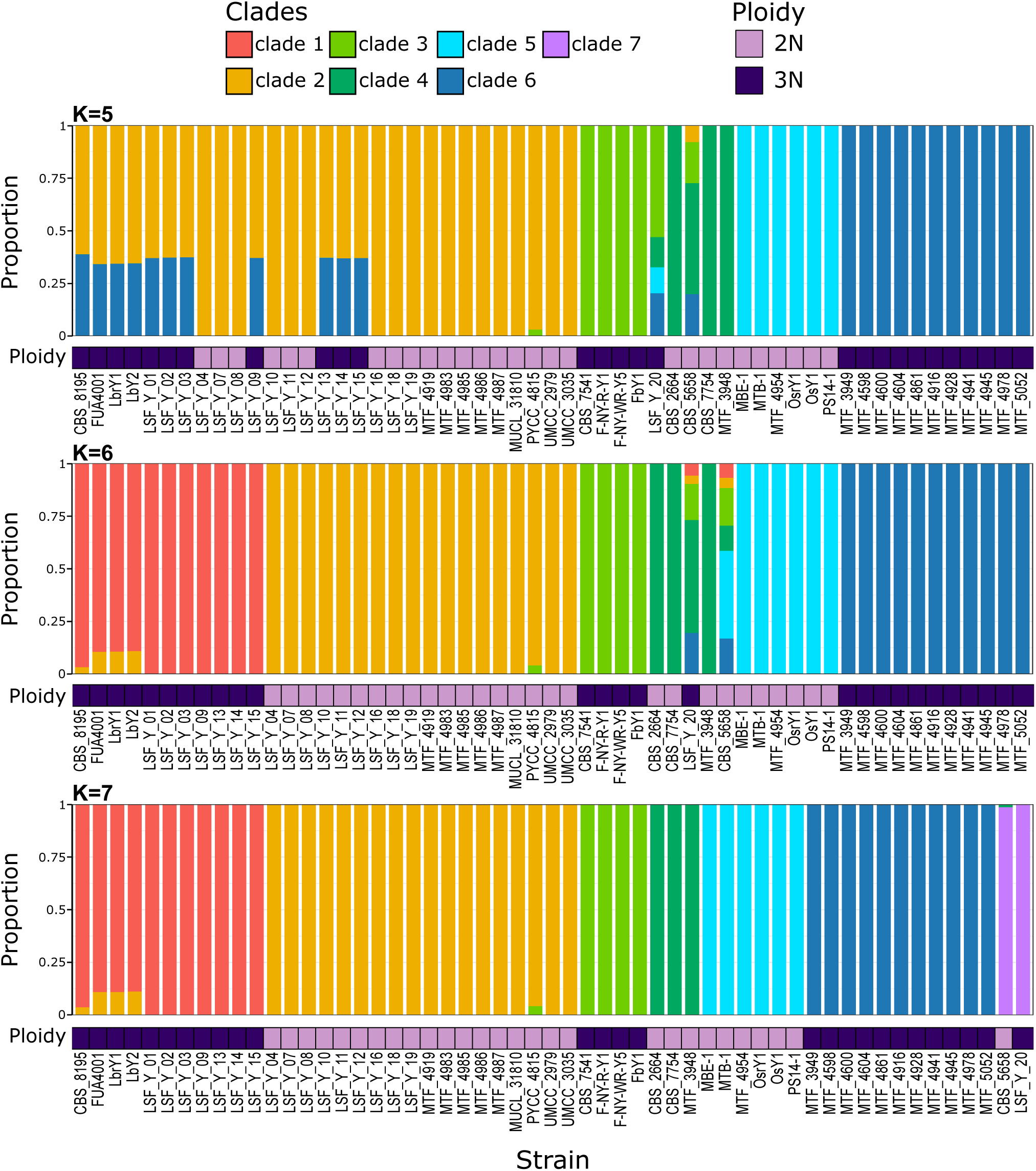
Population structure inferred with ADMIXTURE from the selection of 72,561 SNPs and for a number of subpopulations (K) ranging from five to seven. Columns represent the estimated allocation to subpopulations for each strain. Relative ploidy of strains is provided next to their labels.

Population structure of the 55 strains of *M. humilis* was also investigated using DAPC (Jombart et al. 2010) on the 72,561 biallelic SNPs. According to Bayesian Information Criterion (BIC) values retaining 6 principal component axes (to ensure more than 85% of the variance), the optimal number of clusters was K = 7 (Fig. S2B, Fig. S4). No admixture was predicted for K = 6 to K = 8. Assuming K = 6, the inferred population structure was very similar to that obtained with ADMIXTURE also with K = 6 (Fig. 3 and S3) excepting that the two strains CBS 5658^T^ and LSF_Y_20 were fully associated to clade 4. When considering K = 7, these two latter strains formed a distinct seventh clade, similarly to the prediction of ADMIXTURE for the same value of K. Last, for K = 8, DAPC inference yielded to split clade 1 into two sub-clades (Fig. S3). It is worth noting that one of these sub-clades (clade 1.2) corresponded exactly to the four strains that exhibited moderate admixture in clade 1 in ADMIXTURE predictions (Fig. 3 and S3).

To analyse the genetic distances between strains, a phylogenomic tree was reconstructed using IQ-TREE tool (Minh et al. 2020) based on the 72,561 biallelic SNPs (Fig. 4). The phylogenomic tree was consistent with the six clades previously predicted by the ADMIXTURE and DAPC tools. In addition, the strains whose classification was uncertain, namely CBS 5658^T^ and LSF_Y_20, formed independent branches on the phylogenomic tree, confirming their outlier status among the 55 strains of *M. humilis* (Fig. 4). Therefore, in the remainder of the results, we assume the structuring of the 55 strains of *M. humilis* into six distinct clades and two outliers. Clades 1, 3, and 6 contain triploid strains, with 11, 4 and 11 members respectively. Clades 2, 4 and 5 contain diploid strains, with 18, 3 and 6 members. The diploid strain CBS 5658^T^ will be denoted outlier 1 and the triploid strain LSF_Y_20 will be labeled outlier 2.

**Fig. 4:**
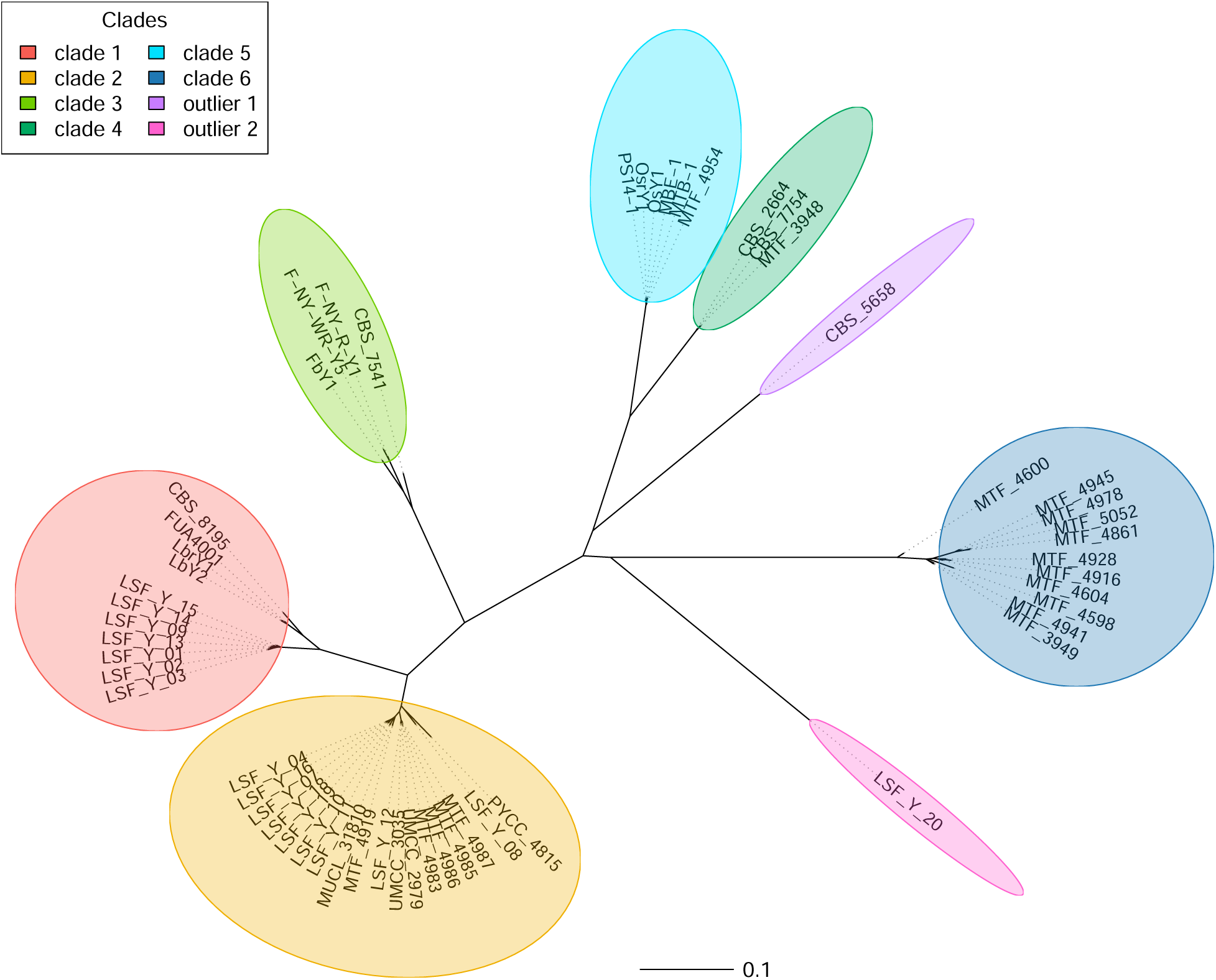
Phylogenomic tree of the 55 *M. humilis* strains, inferred from the selection of 78,561 SNPs. This maximum-likelihood tree is rooted on the TVM+F+R2 model as implemented in IQ-TREE. Colored ellipses correspond to the clades described by ADMIXTURE analysis.

### *M. humilis* lineages demonstrate different patterns of variant accumulation

Investigating SNPs distribution across clades reveals notable differences among clades with the same ploidy. Diploid clades 4 and 5 exhibit equivalent SNPs distribution, with on average 56% and 60% of heterozygous SNPs, respectively. In contrast, clade 2 shows a lower proportion of heterozygous SNPs, at 37% on average (Table S3). Clade differences in genetic diversity are also present among triploid clades, with clades 1 and 6 having on average 71% and 74% heterozygous SNPs, respectively, while clade 3 shows 59% (Table S3).

Looking at LOH proportions and characteristics across clades, we observed substantial differences. The mean number of LOH regions per strain ranged from 29 for the strains of clade 6 to 291 for the strains of clade 2 (Fig. 5 and Table S4), covering on average 683 kb and 5,577 kb, respectively. Despite this important difference, the median size of LOH regions was equivalent between clades. Among all the strains, LOH regions were often found at the ends of contigs (Fig. 5). It is also worth noting that only strains from clade 2 had LOH regions distributed throughout the genome, while for other clades, the distribution of these regions was more scattered. Although a few LOH regions seemed to be shared by all the 55 strains, such as at the end of contig 6, most LOH regions appeared clade-specific (Fig. 5). There were differences in LOH region distribution within the same clade, for example, in clade 1 where strains LbY2, LbrY1, FUA4001 and CBS 8195 had LOH regions that the other strains do not, and conversely. Interestingly, these four strains from clade 1 also stood out from the other seven in the ADMIXTURE analysis (Fig. 3). Lastly, strain CBS 5658^T^ (outlier 1) was the only one strain having LOH that covered entire contigs, namely contigs 7 and 9 (Fig. 5).

**Fig. 5:**
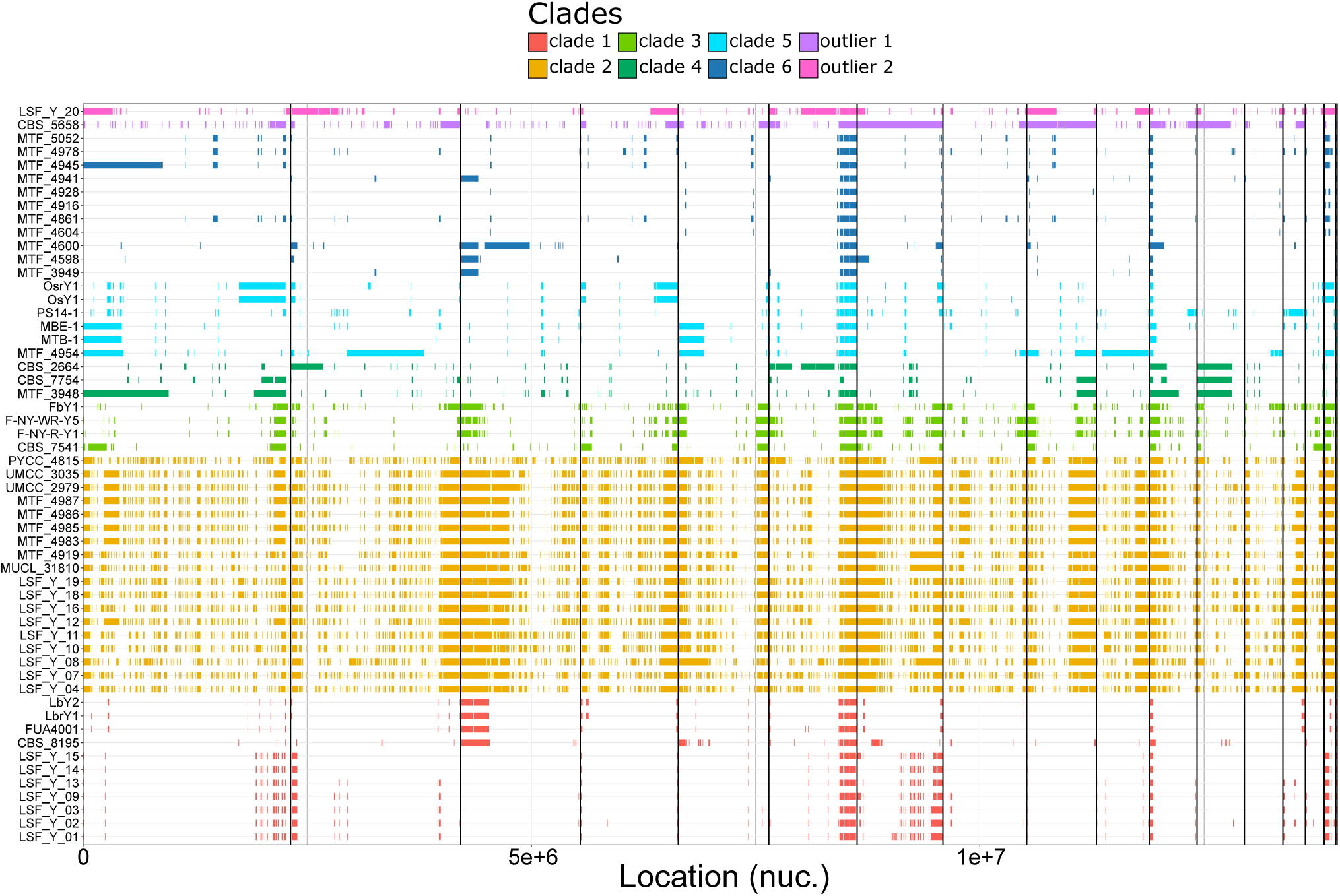
Loss of heterozygosity regions along the genome for each 55 *M. humilis* strains. The 8 colors represent clades from population structure analyses.

### High polymorphism of the ITS region

Using BLASTn search tool (Camacho et al. 2009), we identified 10 unique ITS regions from the draft assemblies of the 55 *M. humilis* strains (Fig. S4). Diploid clades 2 and 4, triploid clade 6, as well as the two outliers had their own unique ITS sequence, while triploid clades 1 and 3 and diploid clade 5 contained multiple distinct ITS sequences (Fig. S4). Interestingly, clade 1 exhibited three distinct ITS sequences suggesting that these triploid strains do not necessarily originate from the same parental strains. To quantify the degree of ITS polymorphism between strains, we aligned the 10 ITS sequences revealing the existence of 26 polymorphic positions out of 563. The most divergent ITS forms were found between the outlier 2 (LSF_Y_20) and two members of clade 5 (OsY1 and OsrY1) with a total of 18 different bases, leading to a sequence identity of 96.80% identity. The ITS sequences of the latter two strains also diverged by 14 bases compared with that of the other four members of clade 5, corresponding to a divergence of over 3%. Fig. S5A summarizes divergence of the different ITS forms when grouped by clade.

### Protein-coding molecular markers suggest a complex origin of clades

Four protein-coding molecular markers were investigated, namely RNA polymerase II largest subunit (RPB1), RNA polymerase II second largest subunit (RPB2), actin (ACT1), and translational elongation factor 1-α (TEF1α). Two paralogous copies of TEF1α were found, disqualifying this gene from further analysis. Genes around TEF1α were found syntenic compared to those of the ohnologous (*i.e.*, paralogous genes that arose from whole genome duplication) copies of TEF1α in *Saccharomyces cerevisiae* (Byrne and Wolfe 2005). The relative divergence of RPB1, RPB2 and ACT1 genes was evaluated when grouped by clade (Fig. S5B, S5C and S5D). Clades and outliers grouped together differently according to the marker considered. Hence, for example, clade 6 grouped with clade 3 when considering RPB1 (Fig. S5B), it was clustered with clade 5 and outlier 1 when comparing RPB2 (Fig. S5C) and it was close to clade 2 when using ACT1 (Fig. S5D). Note that clade 6 was grouped with outlier 1 and clade 4 when considering ITS (Fig. S5A). At the opposite, the clades 1, 2 and 3 always clustered together, whatever the marker considered (Fig. S5). The overall divergence of these three markers was found rather low, less than 2% for RPB1 and RPB2 (Fig. S5B and S5C) and less than 1% for ACT1 (Fig. S5D).

### Strong linkage disequilibrium without decay

Linkage disequilibrium (LD) decay was measured using the PLINK tool (Purcell et al. 2007). LD was estimated as the square of the correlation coefficient between two sites (r^2^). First, this evaluation was done on the 55 strains considering the selection of 72,561 biallelic SNPs. The corresponding LD decay corresponds to the dark grey curve on Fig. 6. With the exception of the very first bases, no LD decay was observed with stable r^2^ values around 0.37 whatever the distance between SNPs. Because LD computation can be impacted by the presence of clones in the compared individuals (Halkett et al. 2005), the same analysis was run considering only one strain per clade, namely MTF 4919 (clade 1), LSF_Y_01 (clade 2), FbY1 (clade 3), MTF 3948 (clade 4), MTF 4954 (clade 5), MTF 4928 (clade 6) and the two outliers strains CBS 5658^T^ and LSF_Y_20. The obtained LD decay, light grey curve on Fig. 6, is very similar to the one taking all strains into account (dark grey curve).

**Fig. 6:**
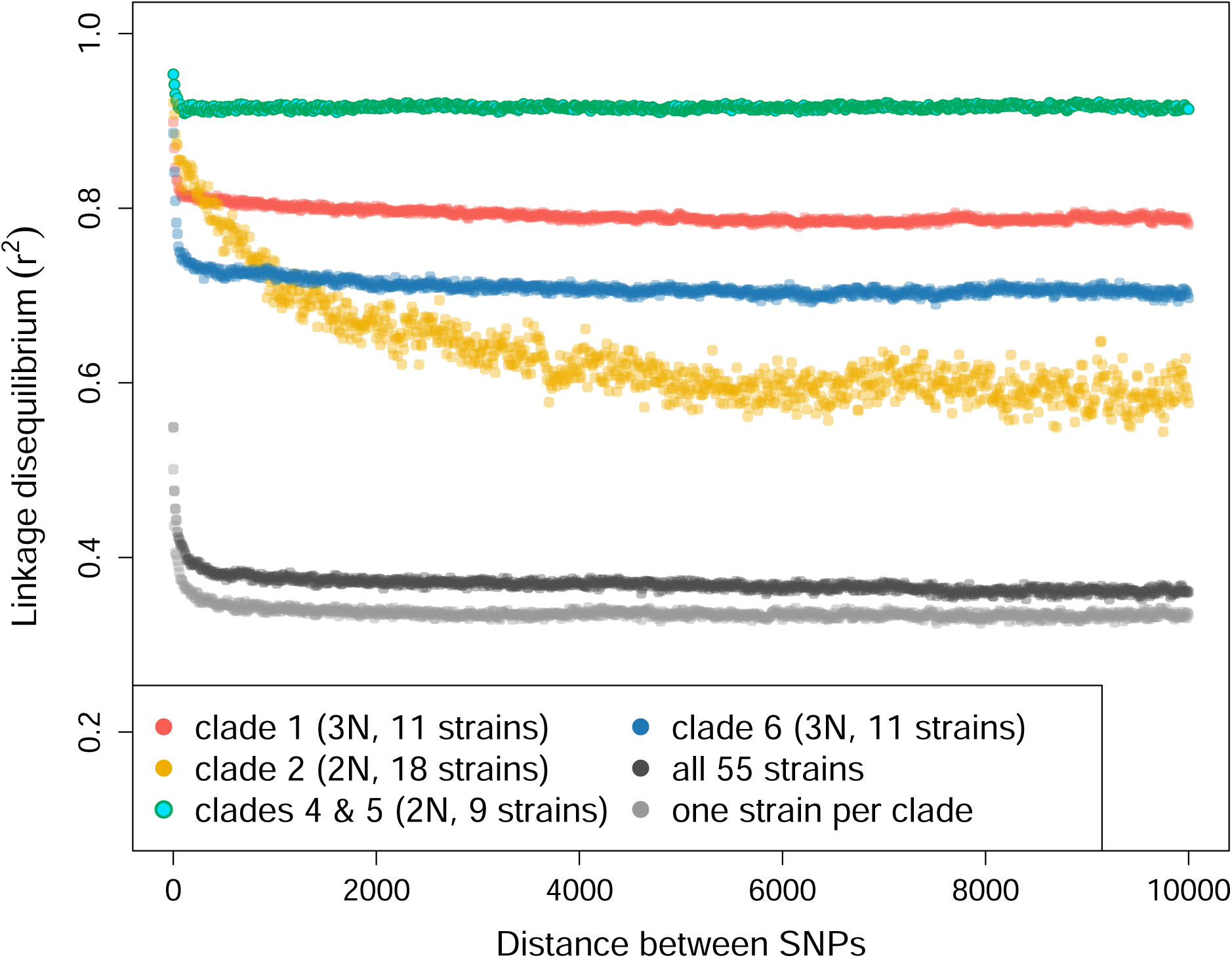
Linkage disequilibrium (LD) decay on different subsets of *M. humilis* strains. For each subset, a selection of biallelic SNPs were performed, which consisted in filtering sites associated to LOHs. For the clades 1, 2 and 6 a total of 1,029,973 SNPs, 225,703 SNPs and 930,931 SNPs were retained respectively. Clade 3 was skipped due to its limited size. Clades 4 and 5 were merged due to their genetic proximity and limited size and yielded a selection of 528,621 SNPs. LD of the whole set of strains was computed on the selection of 72,561 SNPs. The last subset consisted in selecting one strain per clade, including outliers, namely LSF Y 01 (clade 1), MTF 4919 (clade 2), FbY1 (clade 3), MTF 3948 (clade 4), MTF 4954 (clade 5), MTF 4928 (clade 6), CBS 5658T (outlier 1), LSF Y 20 (outlier 2), on the selection of 72,561 SNPs.

To investigate recombination within clades, LD decay was measured clade per clade. Since clades 3, 4 and 5 did not contain enough members to properly compute LD statistics, clade 3 was ignored and clades 4 and 5 were merged for this analysis, given that they both contain diploid strains and are relatively closely related (Fig. 4). To ensure robust results, SNP selections differ between clades as only LOH regions of the corresponding clade were discarded (see details in Materials and Method and in the Fig. 6 caption). Similar to LD decay of the total population, there was no overall LD decay for clades 1, 4/5 and 6 that demonstrated very high average LD values of 0.79, 0.92 and 0.71, respectively. Interestingly, LD statistics of clade 2 were completely different with clear decay from 0 kb to almost 2 kb but still with a high average LD value around 0.61. For the sake of comparison, the intra-clade LD decay analysis was also performed using the set of 72,561 biallelic SNPs (Fig. S6), which yielded similar results.

### All *M. humilis* strains have lost HO endonuclease and silent cassettes

A previous study had shown the absence of the HO gene, which encodes the endonuclease responsible for the mating type switching, in the genome of *M. humilis* strain MAW1 (Mielecki et al. 2024). Investigation of the 55 genomes of *M. humilis* confirmed the absence of the HO gene in all the strains analysed in the present work.

Inspection of the reference genome of *M. humilis* CLIB 1323^T^ allowed the identification of the MAT locus on contig 5 (accession number: CAKOGS010000005.1). It consisted in a complete MATα1 gene (from 859,927 to 860,346) and MATα2 gene (from 859,071 to 859,673, in reverse strand). A relic of the MATa1 gene was also identified downstream of MATα2. Investigation of the surrounding genes showed that the MAT locus of *M. humilis* is syntenic with those of other *Maudiozyma* species and even with the MAT locus of *S. cerevisiae* (Devillers et al. 2022). No silent cassette was detected in the CLIB 1323^T^ assembly. A similar analysis was done on the high quality genome of *M. humilis* strain KH74 (accession number: GCA_037102785.1). An identical MAT locus was found as well as an alternative MAT locus that contained just a complete copy of the MATa1 gene with two introns (accession number: BTGD010000010.1; locations of the 3 exons: 137,810-137,854; 137,915-138,136; 138,201-138,329, in reverse strand). No silent cassette was found in the KH74 assembly.

The complete sequences of the MATa1, MATα1 and MATα2 genes identified above were sought in the draft assemblies of the 55 strains of *M. humilis*. These investigations revealed the presence of MAT loci with both alleles (*i.e.*, MATa1 and MATα1+MATα2) in all the studied strains. We also noted the absence of silent cassettes in the 55 genomes.

### Deciphering genetic origins of triploid strains

Alleles specific to a diploid clade in relation to all the other diploid clades were identified. Specific alleles were retained only if they were homozygous across all the strains from the considered clade. Because they were closely genetically related, strains from clades 4 and 5 were merged and labeled clade 4/5 (Fig. 4). From the set of 72,561 biallelic SNPs, this analysis yielded the identification of 693, 324 and 793 clade-specific homozygous alleles in clade 2, clade 4/5 and outlier 1, respectively. These alleles were then searched for in the triploid strains. Thus, for example, the triploid strain LbrY1 from clade 1 contained 98.4%, 34.9% and 16.4% of the clade-specific alleles of clade 2, clade 4/5 and outlier 1, respectively (Fig. S7). Interestingly, most clade-specific alleles of clade 2 were present in two or three copies in LbrY1 (Fig. S7). These proportions were computed for the 27 triploid strains (Fig. 7A). All the strains from clade 1 contained between 96.1% (strain CBS 8195) and 99.1% (strain LSF_Y_09) of the specific alleles of clade 2 and, on average, only 37.6% and 19.1% of the specific alleles of clade 4/5 and outlier 1, respectively. Triploid strains from clade 3 showed very similar results, notably with 88.3% of the specific alleles of clade 2, on average (Fig. 7A). These proportions were completely different when considering strains from clade 6 whose strains contained on average 37.2%, 65.8% and 49.9% of the specific alleles of clade 2, clade 4/5 and outlier 1, respectively. Last, presence of these specific alleles in the triploid outlier 2 (strain LSF_Y_20) were comparable to those of clade 6 strains (Fig. 7A). In order to determine whether the triploid clades 1 and 3 have a common origin, we first identified all alleles present in these two lineages that were absent from clade 2 among the set of 72,561 biallelic positions. A total of 13,217 and 14,135 alleles were retained for clade 1 and clade 3, respectively. The presence of these alleles was then investigated in all the other lineages, including outliers (Fig. 7B and 7C). Interestingly, 85.5% of clade 1 alleles (11,304/13,217) were found in clade 6 and 25.6% (3,389/13,217) were present only in these two clades (Fig. 7B). Only 9.0% of the retained alleles of clade 1 (1,184/13,217) were not found in any other clade (Fig. 7B). A completely different situation was observed for clade 3, where 38.4% of the retained alleles (5079/14,135) were not found in any other clade and no obvious relationship with another clade was revealed (Fig. 7C). These results demonstrated the occurrence of recent genetic exchanges between diploid clade 2 and triploid clades 1 and 3 as well as between the triploid clades 1 and 6. They also suggested a more ancient relationship between diploid clade 4/5 and triploid clade 6.

**Fig. 7:**
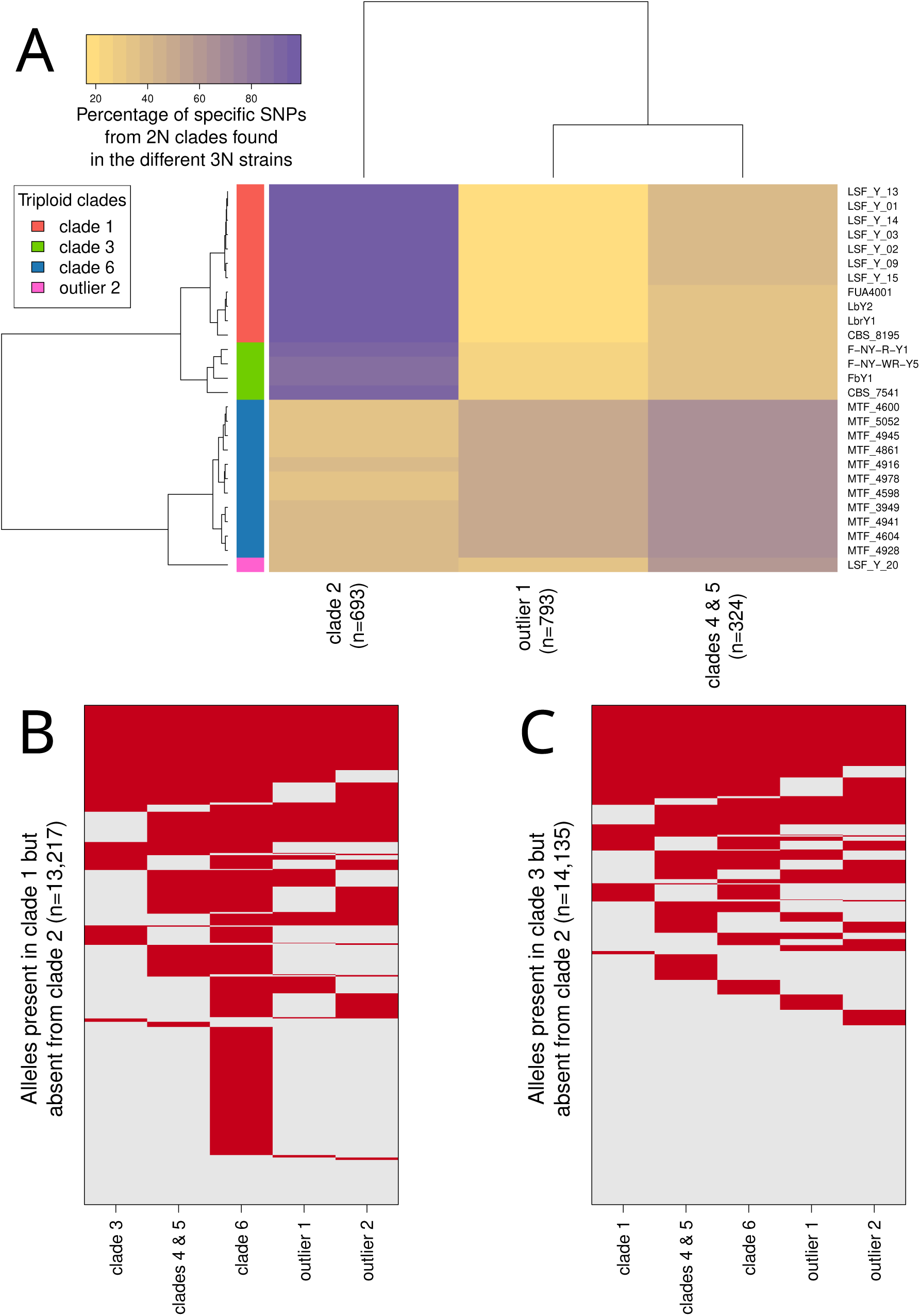
Shared and specific alleles between the *M. humilis* clades. (A) Co-occurrence of homozygous alleles specific to each diploid clade (when considering only diploid clades) with triploid strains. (B) Co-occurrence of alleles from clade 1 that were absent from clade 2 in all the other clades. (C) Co-occurrence of alleles from clade 3 that were absent from clade 2 in all the other clades. Note that for all these analyses, clades 4 and 5 were merged due to their genetic proximity and limited size.

### The phenotypic landscape is associated to the genetic structure

Fermentation performance of the 55 strains of *M. humilis* were evaluated in Sourdough Synthetic Medium (SSM) (Fig. S8). First of all, the absence of block effects was verified by including three biological replicates of the same strain (MTF 4916) in all fermentation runs (Table S5). On average, the latency time was 7.92 ± 0.79 h and the maximum CO_2_ release (Max CO_2_) was 23.1 ± 0.54 g/L. The maximum CO_2_ production rate (Rmax) was 3.06 ± 0.31 g/L/h, and was reached after 13.0 ± 1.58 h of fermentation (tRmax). After 27 h of fermentation, the average viable population density was 1.03 × 10^8^ ± 7.45 × 10^7^ cells/mL with an average mortality rate of 41.1 ± 31.4%.

The two strains identified as outliers on the genomic clustering analysis exhibited different phenotypic characteristics. The strain LSF_Y_20 had fermentation performance and fitness close to the average of all the analysed *M. humilis* strains, while the type strain CBS 5658^T^, showed fermentation performance out of the range of all the analysed *M. humilis*. This latter exhibited the highest latency time (11.31 ± 1.86 h), the longest time to reach the maximum rate of CO2 production (tRmax=23.09 ± 0.45 h), the lowest maximum CO_2_ release (Max CO_2_=21.87 ± 0.64 g/L) and the maximum CO_2_ production rate (Rmax=2.28 ± 0.05 g/L/h) (Fig. S8).

The phenotypic variations of the 53 strains assigned to a clade, excluding the 2 outliers, were mostly driven by the population structure itself (Fig. S8). Significant differences also appeared between diploid and triploid strains for both fitness and fermentation performance (Fig. S9, Table S6). However, a large phenotypic variance was observed within each ploidy (Fig. S9), as illustrated by the fitness difference between clade 2 and clade 4 among diploids (Fig. 8). Indeed, the nested ANOVAs revealed a significant clade effect for each of the 6 phenotypic variables (higher significant p-value < 5 × 10^-5^) (Fig. 8, Table S7).

**Fig. 8:**
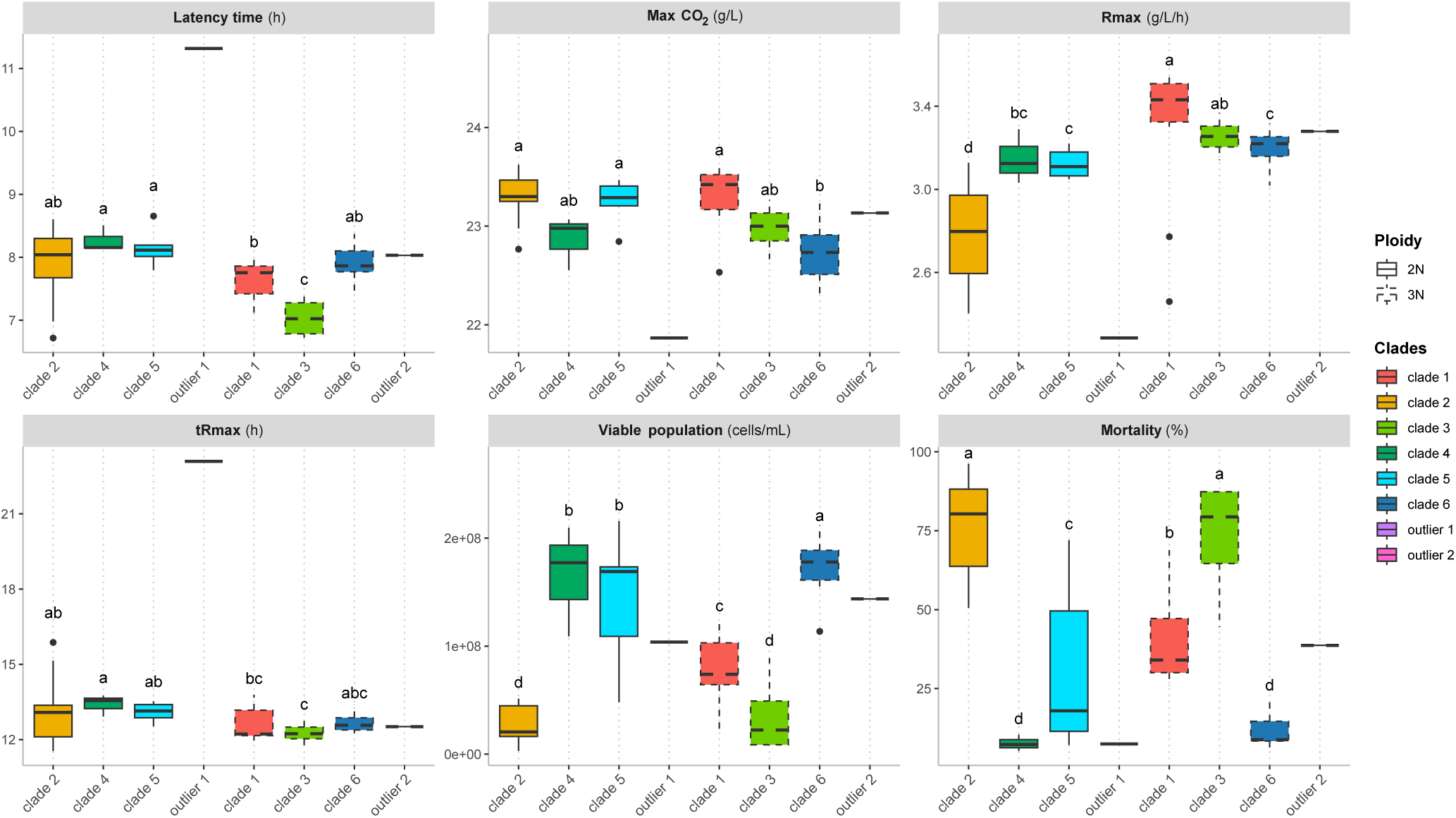
Fermentation performance and fitness in synthetic sourdough medium according to *M. humilis* genetic clades. Fermentation performance was analyzed through four variables: the fermentation latency-phase time (latency time, in h) measured as the time required to reach 1 g/L of *CO*_2_ released, the maximum *CO*_2_ release (max *CO*_2_, in g/L), the maximum *CO*_2_ production rate (Rmax, in g/L/h), and the time to reach the maximum *CO*_2_ production rate (tRmax, in h). Fitness was estimated at 27h of fermentation by the viable population size (viable population, in cells/mL) and the mortality rate (mortality, in %). The outline of the boxplot indicated the ploidy of the strains (dashed line indicates triploids and solid line diploids). A different letter indicated a significant difference based on Tukey’s HSD test.

Clade 3 was the fastest to start fermentation (7.03 ± 0.66 h) and, along with clade 1, had a higher Rmax (3.25 ± 0.14 g/L/h) and lower tRmax (12.30 ± 0.55 h) than the others (Fig. 8). These two groups, which were composed of triploids, also had a lower viable population density and higher mortality rate after 27h of fermentation compared to all the other groups except clade 2, suggesting that they spent more time in a starvation state after the end of fermentation. In addition to low fitness, clade 2 was also characterised by low Rmax (2.78 ± 0.24 g/L/h). All other clades, including diploids and triploids, were intermediate. Clade 6 (22.77 ± 0.57 g/L) presented significantly lower value of maximum CO_2_ release compared to clade 1 (23.29 ± 0.37 g/L), and clade 2 (23.32 ± 0.42 g/L) and 5 (23.25 ± 0.38 g/L) (Fig. 8). Nested ANOVAs also revealed significant differences between strains within the same clade, for each variable, except for maximum CO_2_ release (Table S7, Table S8, Fig. S8).

To synthesize these results, a principal component analysis (PCA) on the 53 strains included in a clade was carried out, including the four fermentation variables (Rmax, tRmax, latency time, Max CO_2_) and two fitness variables (viable population, mortality rate) (Fig. S10). Strains were colored according to their respective clade (Fig. S10A), their country of origin (Fig. S10B) or their substrate of origin (Fig. S10C). The first two axes accounted for 70.9% of the total variation. The first axis (41.2% of the total variation) was explained by variation in viable population and mortality rate and separated clades 2 and 3 from clades 6 and 4 (Fig. S10A and S10D). The second axis (29.7% of the total variation) was mostly explained by the speed of fermentation (latency time and tRmax), and separated clades 3 and 4 (Fig. S10A and S10D). Clade 5 could not be distinguished from any other groups.

A side-by-side representation was used to confront the phylogenomic tree (Fig. 4 and 9, left) with hierarchical clustering based on phenotypic data (Fig. 9, right). A Mantel test revealed a significant positive correlation between the corresponding genotypic and phenotypic matrices (r = 0.3284, p = 0.001), although the coefficient of correlation was relatively low. Inspection of Fig. 9 suggested cases of phenotypic divergence or convergence of strains. In clade 1, the majority of strains were grouped together phenotypically, except for 2 strains (CBS 8195 and FUA4001) that have diverged from the other strains in this clade, but also from each other (Fig. 9). Phenotypic divergence between strains within the same clade was also observed for clade 2. Indeed, 3 strains (LSF_Y_07, LSF_Y_16, LSF_Y_04) diverged phenotypically from the rest of the strains in this group. Interestingly, these 3 strains shared the same geographical origin (Poland) (Fig. 9). Inversely, phenotypic convergence among strains that are phylogenomically distant but originate from the same country, was observed. This was indeed the case of 2 strains (LbrY1 and LbY2) originating from Australia, belonging to clade 1, which clustered on the phenotypic tree with another Australian strain (OsY1) belonging to clade 5 (Fig. 9).

**Fig. 9:**
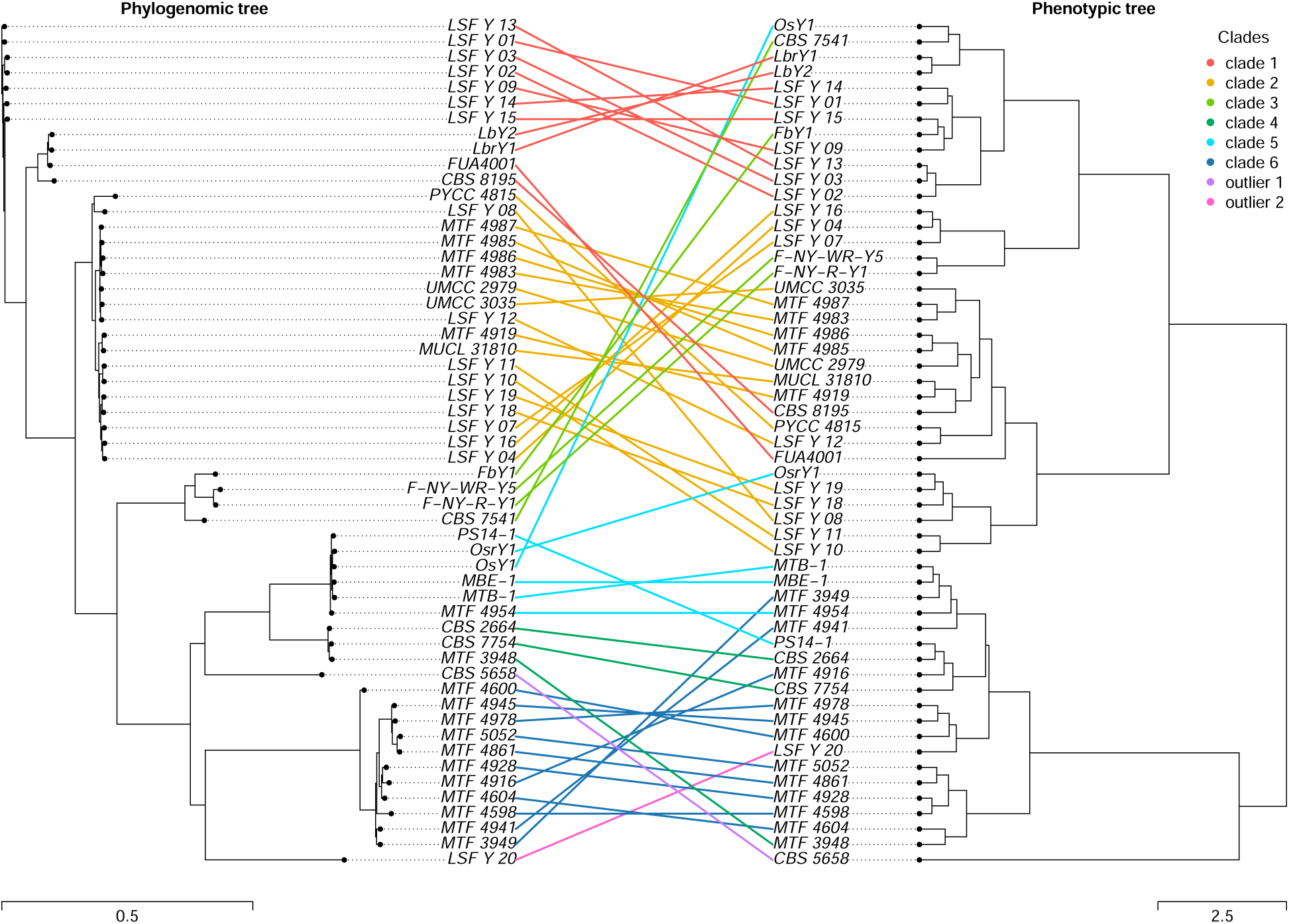
Relationships between the phylogenomic tree (left) and the phenotypic clustering (right). The correspondence between strains is represented by lines colored according to the genetic clade.

### No genetic and phenotypic structuration by geographic and habitat

No evidence of geographic structuration or an effect of the environment of origin was detected. Strains within the same clade originated from different countries, while strains from different clades often shared geographic ranges. Strains originating from rye sourdoughs were distributed over several clades along with wheat sourdough strains suggesting no structure according to the species of cereal used for sourdough preparation. Among the three non-sourdough strains in our dataset, CBS 5658^T^ (from bantu beer) was one of the two outliers, while CBS 7754 (from food dressing) and CBS 2664 (from alpechin) belonged to clade 4, along with the sourdough strain MTF 3948 (Table S1). In addition, no obvious relationships were identified between the ploidy of these strains and their geographical origins and/or their isolation substrate (Table S1).

Similarly, there was no evidence of phenotypic clustering according to geographical (Fig. S10B) or substrate of origin (Fig. S10C). Strains from France, the most represented country, were spread along the first axis. Strains isolated from wheat sourdoughs were distributed all over the plot, as were strains isolated from rye sourdoughs. Therefore no evidence of adaptation to specific geographical area or type of flour was found.

## Discussion

In this study we explored for the first time the evolution of *M. humilis*, the second most frequent yeast species in bread-making. From a collection of 55 strains mainly isolated from sourdough, we characterized six genetic clades and two outlier strains. Genotypic analyses revealed a high heterozygosity in all the studied strains and many genomic structural variations such as ploidy variations or different levels of LOH accumulation. Phenotypic investigation, which included an evaluation of fermentation performance in a synthetic sourdough medium and fitness assessment, demonstrated phenotypic divergence associated with the different genetic lineages.

### *M. humilis* primarily reproduces asexually

Several findings support a predominantly clonal reproduction in *M. humilis*. Observed LD was high and did not decay at all except in clade 2, suggesting limited recombination. This LD pattern is consistent with findings in other yeast species lacking sexual reproduction such as *M. unispora* (Bigey et al. 2025) and *C. albicans* (Ropars et al. 2018). In addition, both DAPC and ADMIXTURE analyses revealed low admixture, further suggesting rare outcrossing events. This restricted admixture may result partly from reproductive isolation between strains of different ploidy levels or with aneuploidy which can disrupt normal meiotic processes, and reduce spore viability, thereby reducing successful mating and genetic exchange (Campbell et al. 1981; Loidl et al. 1994; Albertin et al. 2009; Albertin et al. 2011). The prevalent clonal reproduction of *M. humilis* is also evidenced by the accumulation of LOH regions, a hallmark of clonal populations in which the lack of sexual reproduction prevents the restoration of heterozygosity (Smukowski Heil 2023). Indeed, *M. humilis* harbours a higher overall number of LOH events than in *S. cerevisiae*, with from 15 to 336 LOH regions per strain, compared to 2 to 56 in the *S. cerevisiae* dataset from (Peter et al. 2018).

### The high heterozygosity of *M. humilis* suggests hybridization events

Despite the low level of sexual reproduction, *M. humilis* exhibited a high level of heterozygosity, ranging from 124,231 to 413,957 heterozygous SNPs per genome and 8.88 to 29.59 heterozygous SNPs per kb. Variation of ploidy explains only partly this high level of heterozygosity. While triploids clades had nearly twice as many heterozygous SNPs as diploids (24 vs. 12.7 heterozygous SNPs per kb in average), a high level of heterozygosity in diploids was also found. This level of heterozygosity exceeds the value observed in the worldwide 1,011 strains collection of *S. cerevisiae,* originated from multiple habitats, where the number of heterozygous sites per kb ranged between 0.63 and 6.56 (Peter et al. 2018). Interestingly, the level of heterozygosity in *M. humilis* is comparable to that found in *Candida metapsilosis*, with about 300,000 to 350,000 heterozygous sites per genome (Pryszcz et al. 2015) and *Candida orthopsilosis* with about 100,000 to 400,000 heterozygous sites per genome (Schröder et al. 2016). It has been evaluated that these two species may originate from hybridization of two parental species with an estimated genome divergence of 4.5% and 5%, respectively (Pryszcz et al. 2015; Schröder et al. 2016). Another noteworthy observation is that heterozygosity in *M. humilis* exceeds that of the most heterozygous *Saccharomyces paradoxus* strain WX21, which has an average heterozygosity rate of 0.423% (He et al. 2022). The WX21 strain has been described as arising from multiple intraspecies hybridization events of different lineages whose relative divergence ranged between 0.01 MYA to 0.1 MYA (Eberlein et al. 2019; He et al. 2022). Such a high level of heterozygosity in *M. humilis* genomes strongly suggests that this species originates from the hybridization of quite divergent lineages.

Analysis of taxonomic markers, and more especially the ITS region, further supports strong divergence between the clades and their parental lineages, stronger than what is considered different species in yeasts. Indeed, the ITS variability between clades reaches 96.80% identity (18 polymorphic sites), which is below the commonly used threshold to distinguish Ascomycota yeast species of 98.31 % for ITS (Vu et al. 2016). Such polymorphism on ITS regions have also been observed on hybrid lineages of *C. orthopsilosis* (Sai et al. 2011; Schröder et al. 2016), whose relative divergence parental species was estimated at 5%. These findings reinforce the hypothesis that *M. humilis* evolution implicated multiple hybridization events between distant lineages or even between sister species.

From our knowledge, yeast industries have not constructed *M. humilis* starters through forced hybridization. However, natural hybrids between distinct species spontaneously occurred in yeast creating lineages with increased genetic instability and meiotic infertility (González et al. 2006; Morales and Dujon 2012; Bendixsen et al. 2021; Marsit et al. 2021). All together our results suggest that the strong genetic diversity of *M. humilis* is maintained by rare natural hybridization events, the accumulation of mutations over long evolutionary timescales, eventually a high mutation rate and genome instability, and the persistence of heterozygosity through clonal propagation.

### Multiple transition of ploidy in *M. humilis*

The presence of multiple genetic clades with identical ploidy levels suggests a complex evolutionary history, likely shaped by multiple independent transitions among ploidy states. Such patterns are consistent with well-documented evolutionary process in eukaryotes, where hybridization and polyploidization have recurrently contributed to lineage diversification and phenotypic divergence (Otto and Whitton 2000; Sémon and Wolfe 2007; Akagi et al. 2022).

In yeasts, numerous polyploidization events have been reported both within and between species in the extensively studied genus *Saccharomyces* (Zhu et al. 2016; Duan et al. 2018; Ono et al. 2020; Mozzachiodi and Liti 2022; Tellini et al. 2024). This include allopolyploids populations associated with lager beer (Gibson and Liti 2015), autopolyploids found in ale beer (Gallone et al. 2016; Gonçalves et al. 2016; Fay et al. 2019; Saada et al. 2022), as well as autotetraploids found in bread-making (Albertin et al. 2009; Duan et al. 2018; Peter et al. 2018; Bigey et al. 2021). While the evolutionary consequence of polyploidy has been extensively explored in *Saccharomyces*, their prevalence and significance in other yeast genera remain far less documented. Our results revealed that polyploidization is also common in a closely related genus, which, like *Saccharomyces*, is frequently associated with fermented products.

### Evolutionary trajectories of *M. humilis*

On the basis of 72,561 SNPs and three different methods (ADMIXTURE, DAPC and phylogenetic tree), strain genomes clustered into six genetically distinct clades, comprising three triploid and three diploid lineages, in addition to two genetically isolated strains (CBS 5658^T^ and LSF_Y_20). The combined analysis of the phylogenomic tree, patterns of LOH, shared alleles, ploidy levels, and taxonomic marker sequences allowed us to suggest evolutionary trajectories for the different *M. humilis* clades, including more or less ancient outcrossing events. First, the strong genetic relationship between triploid clade 1 and diploid clade 2 supports the hypothesis that these clades share a recent common ancestor, with triploid clade 1 likely originating from a hybridization event involving an ancestor of clade 2 and an ancestor of clade 6 (Fig. 7A and 7B). Second, clade 3 appears to share ancestry with clades 1 and 2, although this likely occurred earlier, and may originate from a hybridization event involving a lineage absent from our study (Fig. 7A and 7C). Third, the diploid clades 4 and 5 seem to derive from a common ancestral lineage with triploid clade 6 and outlier 2 but in a more distant way than clade 2 when compared to clades 1 and 3. Finally, the diploid outlier 1, which is the type strain of *M. humilis*, may share ancestry with clades 4 and 5, but it remains rather divergent from all the lineages identified in the present work.

Distribution of LOH were found homogenous within *M. humilis* clades (Fig. 5 and Table S4) but differ between clades further confirming the evolutionary trajectories. Each clade carries both large LOH regions (>10 kb), often located at chromosome boundaries generally originating from reciprocal interhomolog crossover, and smaller interstitial LOH fragments generally corresponding to gene conversions (Nguyen et al. 2020; Smukowski Heil 2023). Our results revealed no clear differences in LOH accumulation between diploid and triploid clades but rather differences between clades. Indeed, the average number of LOH blocks for diploid clades identified ranged between 55.2 (clade 5) and 290.6 (clade 2) while for triploid clades it ranged between 29.2 (clade 6) and 121.3 (clade 3) (Table S4). Our findings echo a similar situation observed in a study of diploid hybrid isolates of *Candida orthopsilosis* (Schröder et al. 2016), in which most of the LOH events were found lineages specific. Interestingly, Schroder et al. used the differential accumulation of LOH between clades as a proxy for the relative age of hybridization events (Schröder et al. 2016), revealing that *C. orthopsilosis* may originate from at least 4 distinct and independent hybridization events. Although we are in a similar situation with *M. humilis*, it is more difficult for us to accurately trace the various events that gave rise to the characterized clades, particularly because, unlike *C. orthopsilosis*, no putative parental species that could be the origin of *M. humilis* have been identified. Nevertheless, the diversity of LOH abundance, which correlates with genetic lineages of *M. humilis*, is in line with several studies that have stressed the importance of LOH events in the evolution of the genomes of hybrid and/or asexual species (Smukowski Heil et al. 2017; Lancaster et al. 2019; Bautista et al. 2024).

### Geographical structuration does not explain genetic and phenotypic clustering

Except for clades 1 and 2, only limited admixture was detected among clades suggesting that sourdough strains from different clades rarely meet. Consistent with this hypothesis, strains isolated from the same bakery (Lb, Os, PDC, B17, Table S1) systematically clustered together in the same clade suggesting that strains belonging to different clades rarely co-occurred within a given bakery. This result contrasts with some previous studies of *S. cerevisiae* populations, in which higher levels of mixed ancestry and outcrossing have been associated with anthropogenic activities (Magwene et al. 2011; Wang et al. 2012; Cromie et al. 2013; Legras et al. 2018; Fay et al. 2019).

Nevertheless, long distance migration appears to occur. Indeed, each clade includes strains coming from diverse countries. This lack of geographical structuration and isolation by distance suggests human-mediated long-distance dispersal, rather than local dispersion. Similar patterns have been observed in other yeast species, notably *S. cerevisiae*, in which genetic differentiation is more strongly driven by ecological and functional specialization than by geographic origin (Legras et al. 2007; Liti et al. 2009; Sicard and Legras 2011; Legras et al. 2018; Peter et al. 2018) The vectors responsible for the dispersal of sourdough yeasts remain to be elucidated and may differ among species and clades. Study based on extensive sampling of flours on-farm milling or artisanal millers, together with their associated sourdoughs, failed to detect *S. cerevisiae* or *M. humilis* in 44 flours, suggesting that flour is unlikely a major dispersal vector for sourdough yeast species (von Gastrow, Michel, et al. 2023). In contrast, two other studies detected both *Maudiozyma* and *Saccharomyces* genus in some commercial flours, suggesting that industrial flour production may contribute to the dispersal of *Maudiozyma* (Minervini et al. 2015; Taheri et al. 2026). In addition, *S. cerevisiae* and *M. humilis* have been detected on human hands (Reese et al. 2020), pointing to a potential role of human-mediated transmission in shaping yeast distribution. However, *M. humilis* has so far not been detected on humans. Together, these observations highlight the need for further environmental genomics investigations of habitats associated with bakery environments in order to identify dispersal routes and assess the relative contributions of different vectors to yeast species evolution.

### Historical contingency rather than ploidy drives phenotypic variation in *M. humilis*

Although increased ploidy is often assumed to confer fitness advantages in domesticated plant and fungal populations, our results reveal a more complex relationship between phenotype and genotype. The six genetic clades showed phenotypic divergence and we detected a significant correlation between phenotypic and genomic distances among pairwise strains. However, this association was not related to ploidy variation alone. Indeed, triploid strains from clades 1 and 3 showed significantly lower fitness in average than diploid strains of clades 4 and 5, challenging the assumption of a systematic fitness advantage associated with higher ploidy.

Polyploidy is expected to confer fitness advantages by masking recessive deleterious mutations (Otto and Whitton 2000; Fischer et al. 2021), and by increasing the likelihood of the emergence of beneficial mutations, of novel allelic interactions, or by creating genetic and epigenetic rearrangements (Gerstein and Otto 2009; Akagi et al. 2022). Consistent with this view, several studies have documented the fitness and fermentation advantages of polyploid yeast populations in human-maintained environments. For example, Krogerus *et al*. showed that *S. cerevisiae* polyploids hybrids exhibited increased stress tolerance, and higher fermentation performance to their parental strains (Krogerus et al. 2016). In contrast, our findings suggest that historical contingency and lineage-specific evolutionary trajectories may have a stronger influence on fitness than ploidy level in *M. humilis*. A similar pattern has been reported in bread-making populations of *S. cerevisiae*, in which tetraploidy does not confer a fitness advantage. Although commercial tetraploid *S. cerevisiae* strains exhibit a shorter onset of fermentation, diploid sourdough strains display higher fitness in sourdough mimicking medium (Bigey et al. 2021).

Hybridization is known as a source of variation for adaptation, facilitating phenotypic divergence and adaptive radiation (Lewontin and Birch 1966; Kagawa and Takimoto 2018). However, several studies have shown that hybridization may have opposite effects depending on the life history traits leading to the persistence of parental and hybrid lineages (Clo et al. 2021; Rosser et al. 2024). Variation of fitness among *M. humilis* clades indicates that selective pressures may have differed across lineages leading to phenotypic divergence potentially reflecting differences in colonization history within sourdough or differences in the intensity of bakery practices. Diploid clades 4 and 5, as well as triploid clade 6 exhibited significantly higher fitness than the other clades, suggesting stronger natural selection and/or a greater number of generations in the sourdough environment. In contrast, triploid clades 1 and 3 showed a relatively high fermentation performance, characterized by short latency and high Rmax, with lower overall fitness, a pattern consistent with human-mediated selection favoring fermentation efficiency for intensive baking practices over population growth and survival. Interestingly, triploid clade 6 combined both high fitness and strong fermentation performance, indicating that there is no evidence of trade-off between these traits. Finally, clade 2 displayed the lowest fitness and fermentation performance across all clades, which may reflect a more recent colonization of the sourdough environment. Alternatively, the low level of heterozygosity and high level of LOH present in this clade may reduce its potential for adaptation in the sourdough environment.

### *M. humilis*, a domesticated species?

A wide variety of hallmarks have been described in the genome of domesticated species, such as high heterozygosity, polyploidy, aneuploidy, loss of heterozygosity (LOH), chromosomal rearrangements, or hybridization (Legras et al. 2014; Gallone et al. 2016; D’Angiolo et al. 2020; De Chiara et al. 2022). Most of these signatures have been identified in the investigated genomes of *M. humilis*. Domestication may also impact yeast life-cycle (Fischer et al. 2021; De Chiara et al. 2022; Becerra-Rodríguez et al. 2026). Our results reveal that *M. humilis* reproduces predominantly asexually, and is heterothallic, with a total deletion of the HO gene and the silent cassettes. Loss of the ability to perform mating-type switching, either through loss-of-function mutation of *HO* endonuclease or the loss of silent cassettes, has been found in several *Saccharomyces* domesticated populations and can be seen as an evolutionary strategy favouring external outcrossing if a sexual cycle is initiated (Becerra-Rodriguez et al. 2026). These observations suggest convergent evolution of heterothallism associated with domestication across yeast genus.

All together our data reveal the complex evolution of sourdough *M. humilis* populations, involving hybridization, polyploidy, dynamic LOH, the evolution of life history traits and divergent fermentation ability and fitness. So far *M. humilis* has been mostly isolated from fermented products limiting the analysis of the impact of domestication. However, extending this study to strains from other fermented products will shed light on the role of adaptation in the evolution of *M. humilis*.

## Materials and Methods

### Strain collection

This study was performed using a collection of 55 *M. humilis* strains chosen to represent diverse ecological and geographical origins. Each strain has been derived from a single colony. Forty two strains were isolated from Europe, and the other strains originated from South Africa (1), Australia (5), Canada (1), China (2), Lebanon (1), Morocco (1) and USA (2). Among the 55 strains, 52 were retrieved from bread-related environments and 3 from diverse ecological origins, namely food dressing, bantu beer and alpechin (liquid waste from olive oil mills) (Table S1). These strains were obtained from public collections (CBS, CIRM-Levures, MUCL, UMCC, PYCC), and from several collaborators. The Spanish strains were kindly provided by Dr. Mercedes Tamame (Universidad de Salamanca, (Chiva et al. 2020)), the USA strains by Pr. Luc de Vuyst (Vrije Universiteit Brussel, (Comasio et al. 2020)), the Canadian strain by Pr. Michael Gänzle (University of Alberta), the Lebanese strain by Dr. Pamela Bechara and Pr. Marie-José Ayoub (Lebanese University, (Bechara et al. 2025)), and the Australian strains from Dr. Anna Wittwer and Pr. Kate Howell (University of Melbourne, (Wittwer et al. 2026)). French strains were isolated from sourdoughs collected from different bakers and farmer-bakers (associated references in Table S1). Finally, 17 strains were provided by a private supplier (LESAFFRE, Table S1). The Access and Benefit-sharing of all these strains follow the rule of each country and respect the provider demand. All strains were stored at -80 °C in YPD supplemented with 15% (v/v) glycerol.

### Genome sequencing

All *M. humilis* strains were subjected to Illumina paired-end sequencing. Total genomic DNA was extracted using a method adapted from Hoffman and Winston 1987. Yeast cell cultures were grown overnight at 30 °C in 50 mL of YPD medium and then harvested by centrifugation (10,000 g, 2 min). Cells were washed in 1 mL water and collected by centrifugation (10,000 g, 1 min), resuspended in 300 µL of lysis buffer (Tris 10 mM pH8, EDTA 1 mM, NaCl 100 mM, Triton 2%, SDS 1%) with glass beads (425-600 µm), as well as 300 µL of phenol/chloroform/isoamyl alcohol 25:24:1. Tubes were vortexed for 5 min, cooled on ice for 5 min and vortexed for 5 min. Solutions were centrifuged (10,000 g, 5 min), supernatants were collected and 800 µL of pure ethanol were added and mixed gently. After brief centrifugation (5,000 g, 1 min), the DNA pellet was washed with 70% ethanol, re-suspended within 100 µL of Tris-EDTA, and kept at 4 °C overnight. One microliter of RNase A (10 mg/mL) was added to the samples and incubated in a dry bath at 37 °C for 30 min, then stored at 4 °C.

Genomic Illumina sequencing libraries were prepared and subjected to paired-end sequencing (2×150 bp) on Illumina NovaSeq 6000 at GeT-PlaGe Toulouse INRAE or Eurofins.

### Read trimming and draft genome assembly

Illumina paired-end reads were cleaned with fastp tool (v0.23.2) (Chen 2023) to trim low quality regions, technical adaptors and poly-G monomers. Only reads greater than 50 pb after trimming have been retained. A draft genome assembly of each sample was generated using SPAdes tool (v3.15.5) (Prjibelski et al. 2020) with default parameter values.

### Read mapping and variant calling

For each strain, trimmed Illumina paired-end reads were mapped against the reference *M. humilis* CLIB 1323^T^ genome (14.22 × 10^6^ bp, 16 contigs, accession number : GCA_933934105.1) with BWA mem (v0.7.17 default parameters) (Liti et al. 2009). Because the mitochondrial chromosome was missing from this assembly, short reads of the reference genome strain (CLIB 1323^T^, SRA: ERR5775463) were reassembled using SPAdes (v3.15.5) and the identified mitochondrial sequence was retrieved to complement the reference genome used. This mitochondrial chromosome is available on the following data repository: https://doi.org/10.5281/zenodo.18642067. Alignments were post-processed using SAMtools fixmate, SAMtools sort and SAMtools markdup (1.15.1) (Danecek et al. 2021). Coverage was retrieved for each position with SAMtools depth function and mean coverage values along the genome were plotted using 5 kb and 1 kb step sliding windows. Single-nucleotide polymorphisms (SNPs) and insertions/deletions (INDELs) were identified using the HaplotypeCaller function in GATK version 4.5.0.0 (McKenna et al. 2010), with ploidy set to 2. For the global analysis, we combined the variant files into a single file using the CombineGVCFs function in GATK. We then extracted only the variant positions using the GenotypeGVCFs function. Biallelic and multiallelic SNPs were separated using the SelectVariants function in GATK for distinct analyses. SNPs were filtered using GATK VariantFiltration with these following parameters: QD < 2.0, LowQD, ReadPosRankSum < −8.0, ReadPosRankSum > 8.0, FS > 60.0, HightFS, MQRankSum < −2.5, MQRankSum > 2.5, MQ < 40.0, Low_MQ, SOR > 3.0, High_SOR, Bad_MQRS, Bad_RPRS ; and SelectVariants function, following the hard-filtering recommendations for germline short variants from the GATK documentation (De Summa et al. 2017). Finally, we used GATK VariantsToTable to extract genotype and allele frequencies from the filtered vcf file. Allele frequencies, depth and number of heterozygous sites were then computed along the genome using R (R version 4.3.1 and Rstudio 2024.4.2.764).

### Ploidy assessment

Strain ploidy was analysed by flow cytometry as described in (Bigey et al. 2021). Flow cytometry profiles were compared to those obtained for a closely related species, *Monosporozyma unispora*, formerly *Kazachstania unispora* (Bigey et al. 2025). The three *M. unispora* strains CLIB 234 (haploid), MUCL 45647 (diploid) and MTF 4423 (triploid) were used as references. In addition, ploidy were confirmed based on allele frequency distribution at heterozygous SNPs within each sample. They were considered diploids when the distribution was centered on 0.5 and triploids if the distribution included frequencies peaks at 0.33 and 0.66.

### Detection of regions with loss of heterozygosity

Identification of regions with loss of heterozygosity (LOH) was carried out with JLOH v1.1.0 (Schiavinato et al. 2023) with parameters “--min-snps-kbp” set to 1.1, “--min-length” to 2000, “--min-af” to 0.13 and “--max-af” to 0.87. SNPs detected inside LOH blocks were then removed from the filtered vcf file for the analysis of population structure.

To characterize LOH among our strains, we applied a filtering process to retain only high-confidence LOH blocks. We selected blocks with coverage exceeding 60% of the mean coverage, merged adjacent LOH regions separated by fewer than 0.3% heterozygous SNPs, and excluded blocks shorter than 3,000 bp.

### Linkage disequilibrium

Linkage disequilibrium (LD) was computed as r^2^, the square of the correlation coefficient between pairs of loci, with PLINK v1.90 (Purcell et al. 2007). Pairs of SNPs were sampled on 10 kb windows (--ld-window-kb 10) and 1,000 SNPs windows size (--ld-window 1000) and excluding pairs of SNPs with a r^2^ statistics below 0.1 (--ld-window-r2 0.1). To limit bias, LD statistics were computed on selections of SNPs excluding LOH regions.

### Population structure

Population structure was inferred using model-based clustering algorithm and a discriminant analysis of principal components (DAPC), on the same selection of SNPs as for the phylogenomic analysis. The Bayesian clustering analysis was carried out using ADMIXTURE v1.3.0 (Alexander et al. 2009). Using between 1 and 10 populations for the 55 strains, the optimal K value was selected based on the lowest cross-validation error. The DAPC analysis was conducted using the R package adegenet (Jombart et al. 2010), and the optimal number of clusters was determined using the Bayesian Information Criterion (BIC).

### Phylogenomic analysis

IQ-TREE (v2.0.7) (Minh et al. 2020) with 1000 ultrafast bootstrap replicates was used to infer phylogenomic relationships between the 55 *M. humilis* using a selection of SNPs chosen outside regions with LOH. The model finder procedure (-m MFP) was used to select the best model to infer the phylogenetics tree. As our dataset does not include ancestral strain or any previous datation, we used an unrooted tree without defined outgroup.

### Taxonomic molecular markers

ITS regions of the different *M. humilis* strain were retrieved from their draft assemblies using BLASTn search tool (Camacho et al. 2009) and the ITS sequence of type strain of *M. humilis* NRRL Y-17074^T^ (GenBank accession: AY046174.1). Each unique ITS identified among the 55 strains was used as a BLASTn query to classify the strains based on their ITS sequence. The ITS clusters were then compared to clades to assess their correspondence. These ITS sequences were then aligned with BioEdit v7.7.1 (Hall 1999) to identify any polymorphism.

As an alternative or complement to rDNA based markers like ITS, markers derived from protein-coding genes are also often used for taxonomic and phylogeny purposes. These are generally essential genes, present in a single copy in the genome. Four commonly used markers for yeast taxonomic studies were considered, namely actin (ACT1), RNA polymerase II largest subunit (RPB1), RNA polymerase II second largest subunit (RPB2) and translational elongation factor 1-α (TEF1α). These four genes were first searched in the reference genome, strain CLIB 1323^T^. Then, using the variant calling data, the corresponding genotypes of the four markers for the 55 strains were determined. Polymorphic sites were encoded with the IUPAC ambiguity code. Resulting sequences were then analyzed with the R packages bioseq (v0.1.5) (Keck 2020).

### Detection of the HO gene

The presence of the HO gene that encodes for the endonuclease responsible for the mating-type switching, was investigated using a tBLASTn (v2.15.0+) (Camacho et al. 2009) search against all the draft genome assemblies of all compared strains. The HO protein sequence of *Maudiozyma bulderi* CLIB 596 (GenBank accession: CAL1764204.1) was used as a query.

### Searching for mating-type genes

Mating-type gene were first searched in the reference genomes of *M. humilis* CLIB 1323^T^ (GenBank accession: GCA_933934105.1) and KH74 (GenBank accession: GCA_037102785.1). Because no annotation was available for these genes in that species, mating-type genes from a closely related species, *Maudiozyma barnetti* (Devillers et al. 2022), were used as query in tBLASTn searches (v2.15.0+) (Camacho et al. 2009). Candidate genes were then used to identify all putative loci related to mating-type in the genomes of the 55 strains of *M. humilis*.

### Sourdough Synthetic Medium

Fermentations were performed in Sourdough Synthetic Medium (SSM) adapted from (Bigey et al. 2021). SSM contained at final concentrations 24 g/L wheat peptone (Sigma), 200 mg/L MgSO_4_.7H_2_O (Fluka), 50 mg/L MnSO_4_.H_2_O (Sigma), 4 g/L KH_2_PO_4_ (Sigma) and 4 g/L K_2_HPO_4_ (Sigma), adjusted at pH = 4.5 with citric acid. The sterilized medium was completed with different solutions to reach at final concentration (i) 50 g/L glucose (solution filtered at 0.22 µm) (Panreac Applichem) (ii) 1.4 g/L linoleic acid (Sigma), 1 mL Tween 80 (Sigma) and 98 mg/L β-sitosterol (Sigma) (Melis and Delcour 2020) and (iii) 0.2 mg/L of each vitamin (cobalamin, folic acid, nicotinic acid, pantothenic acid, pyridoxal-phosphate, thiamine (Sigma)).

### Assessing fermentation performance

Fermentations were carried out as described in Boudaoud et al. 2021. Strains were isolated on YPD plates from frozen stocks (stored at -80 °C), and incubated at 28 °C for 20 h. Then, one colony was inoculated in 3 mL of YPD liquid to prepare precultures, incubated at 28 °C for 20 h at 220 rpm. After centrifugation at 4,256 g for 5 min of precultures, cells were suspended in 1 mL of SSM. Cell concentration was measured with an Attune NxT^TM^ flow cytometer (Invitrogen by Thermo Fisher Scientific) equipped with an CytKick^TM^ Max Autosampler (Invitrogen by Thermo Fisher Scientific). Cell concentration was adjusted at 10^6^ cells/mL in 15 mL of SSM into a glass tube equipped with a filter tip to allow CO_2_ release. Fermentations were carried out over 27 h at 24 °C with constant stirring (350 rpm). An automated robotic system (PlateButler® Robotic System by Lab Services) (Bloem et al. 2018) was used and allowed online monitoring of the fermentation by measuring every 50 min for each sample, the weight loss which is correlated to CO_2_ release. Data was automatically inserted into the ALFIS (Alcoholic Fermentation Information System) software. Based on polynomial smoothing, fermentation parameters were estimated from the CO_2_ accumulation curve over time (Sablayrolles et al. 1987; Michel et al. 2023): the fermentation latency-phase time (latency time, in h) corresponding to the time required to reach 1 g/L of CO_2_ released, the maximum CO_2_ release (max CO_2_, in g/L), the maximum CO_2_ production rate (Rmax, in g/L/h) and the time to reach the maximum CO_2_ production rate (tRmax, in h). At least three independent experimentations were carried out for each strain. All trials were randomly distributed over 7 fermentation runs. Three replicates of the MTF 4916 strain were placed in all fermentation runs to verify the absence of block effect.

### Assessing fitness

After 27 h of fermentation, samples were collected to determine the mortality rate and the viable population. Fifty µL of each sample were centrifuged (3,363 g for 5 min at 4 °C) and diluted in Phosphate Buffered Saline (PBS, Sigma). Samples were labeled with 1 µg/mL propidium iodide (PI, Calbiochem), incubated 10 min in the dark before measurement and analysed with an Attune NxT^TM^ flow cytometer.

### Statistical analysis

Analysis of variance (ANOVA) was performed to evaluate the difference between diploid and triploid strains for each fermentation and fitness variable (latency, max CO_2_, Rmax, tRmax, viable population, mortality). Nested ANOVA were carried out to investigate the variation of each fermentation and fitness variable among clades and among strains within a clade. After each nested ANOVA, multiple comparisons of means according to the Tukey’s HSD test were used to determine (i) for each variable which clades were significantly different from each other and (ii) for each variable (except max CO_2_) which strains were significantly different from each other. Principal component analysis (PCA) was performed on fermentation performance and fitness data of strains belonging to clades (outliers were excluded). All analysis and graphical representations were performed with RStudio (R version 4.3.1 and Rstudio 2024.4.2.764).

### Comparison of phenotypic and phylogenomic trees

The relationship between phylogenomic distance and multivariate phenotypic distance between strains was investigated. For that, based on the 6 phenotypic parameters measured for the 55 strains studied, a phenotypic tree was constructed by successive steps, involving data standardization, creation of a distance matrix (euclidean method), then an hclust object (ward.D2 method) transformed into a tree (as.phylo function from the R package ape (Paradis and Schliep 2019)). The obtained phenotypic based tree was then compared to the phylogenomic tree obtained in section 2.8.1. To do so, we visualized a potential correlation between phenotypic and phylogenomic trees by optimizing vertical matching of tree tips using function cophylo in the R package phytools (Revell 2024).

To better assess the relationship between the phenotypic and phylogenomic trees, their associated distance matrices were compared. It was used (i) for the phenotypic tree, the Euclidean distance on which it is based and (ii) for the phylogenomic tree, the cophenetic distance extracted from the tree using the cophenetic function. The correlation between both distance matrices was tested with Mantel test (permutation = 999, method = spearman) in the R package vegan (Oksanen et al. 2024).

## Supporting information

Supplementary figures from S1 to S10

Supplementary tables from S1 to S8

## Acknowledgements

We thank Anna Wittwer, Marie-José Ayoub, Pierre Abi Nakhoul, Mercedes Tamame, Luc de Vuyst, Michael Gänzle, Lesaffre and bakers for providing strains. We thank Cécile Neuvéglise for ordering strains from the international collection. We thank Jean-Luc Legras for its scientific suggestions and comments. We thank Stéphane Guézenec for his contribution to the DNA extractions and management of the yeast collection.

## Authors contribution

**Conceptualization**: D.Si., D.Se., H.D.; **Methodology**: M.L., J.R., D.Si., D.Se., H.D.; **Software**: J.R., H.D.; **Validation**; M.L., J.R., D.Si., H.D.; **Formal analysis**: M.L., J.R., T.N., H.D.; **Investigation**: M.L., J.R., D.Si., T.M., L.A., D.Se., H.D.; **Resources**: D.Si., K.H., P.B., L.A., D.Se.; **Data Curation**: M.L., J.R., H.D.; **Writing - Original Draft**: M.L., J.R., D.Si., H.D.; **Writing - Review & Editing**: all authors; **Visualization**: M.L., J.R., T.N., H.D.; **Supervision**: D.Si., D.Se., H.D.; **Project administration**: D.Si., H.D.; **Funding acquisition**: D.Si.; All authors read, revised and approved the manuscript.

## Data availability

All sequencing data is available at the EBI/NCBI databases under the project PRJEB89020. The detailed list of accession number for samples and SRA (raw reads) for each strain is provided Table S1. Complementary data, material and scripts used and/or generated in the present work are available at https://doi.org/10.5281/zenodo.18642067.

## Supplementary

**Supplementary_Figures.pdf**: Supplementary figures from Fig. S1 to Fig. S10.

**Supplementary_Tables.xlsx**: Supplementary tables from Table S1 to Table S8.

