## Supplementary figures from S1 to S10 for "Genotypic and phenotypic diversity of *Maudiozyma humilis*: the multiple evolutionary trajectories of a domesticated yeast"

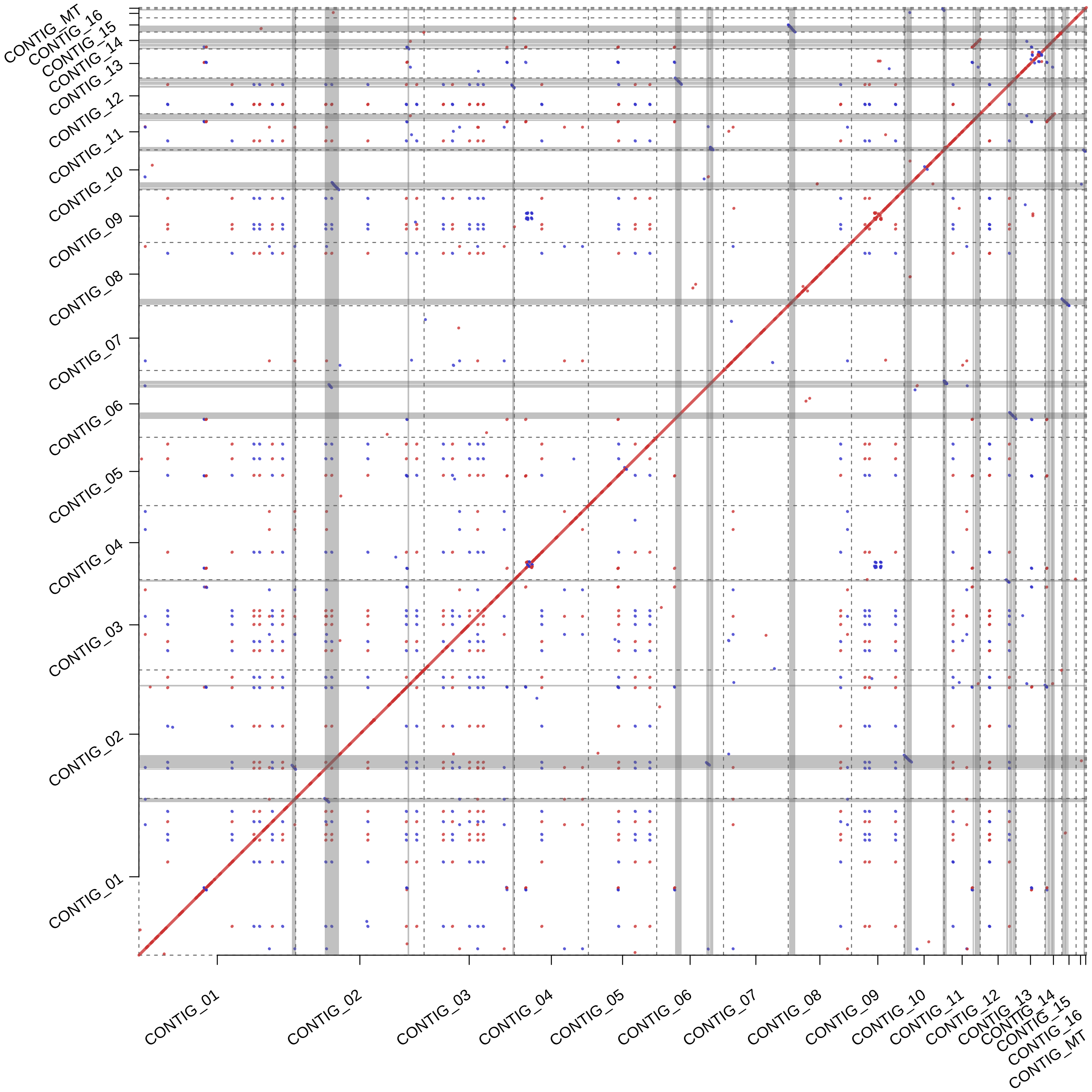


**Fig. S1**: Self-to-self MUMmer analysis of the reference genome used in or analysis (type strain, CLIB $1323^T$). Large redundant regions ($\geq$ 20kb) were identified (shaded regions). They may result from technical issues such as unresolved haplotype during the assembly or may reflect biological features like large transposable elements. They were masked from our subsequent analyses to avoid detections of spurious structural variations or LOH regions.

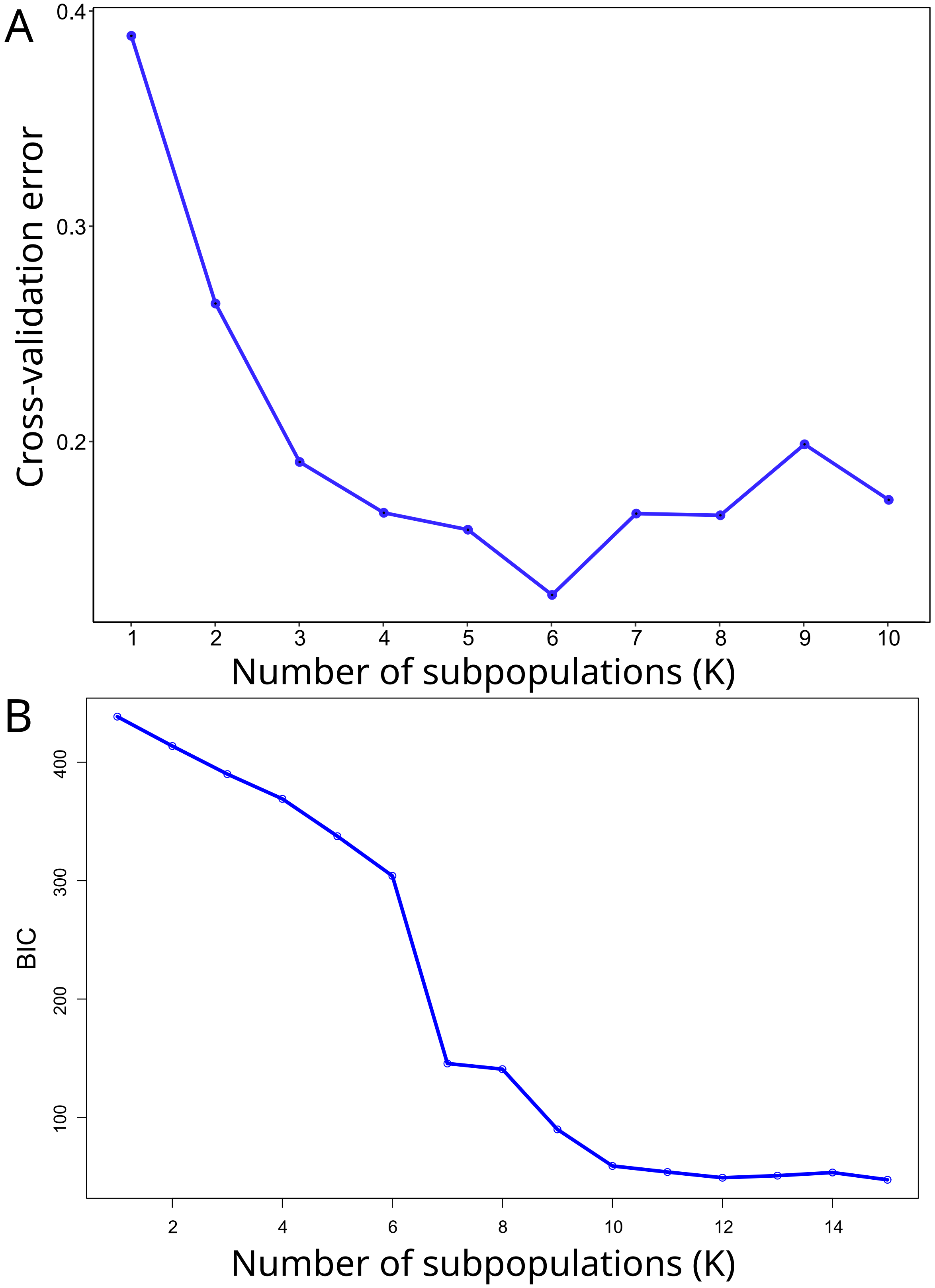


**Fig. S2**: Statistical evaluation of subpopulation size. (A) Plot of ADMIXTURE cross validation error from K = 1 to K = 10. (B) Plot of DAPC BIC values from K = 1 to 15.

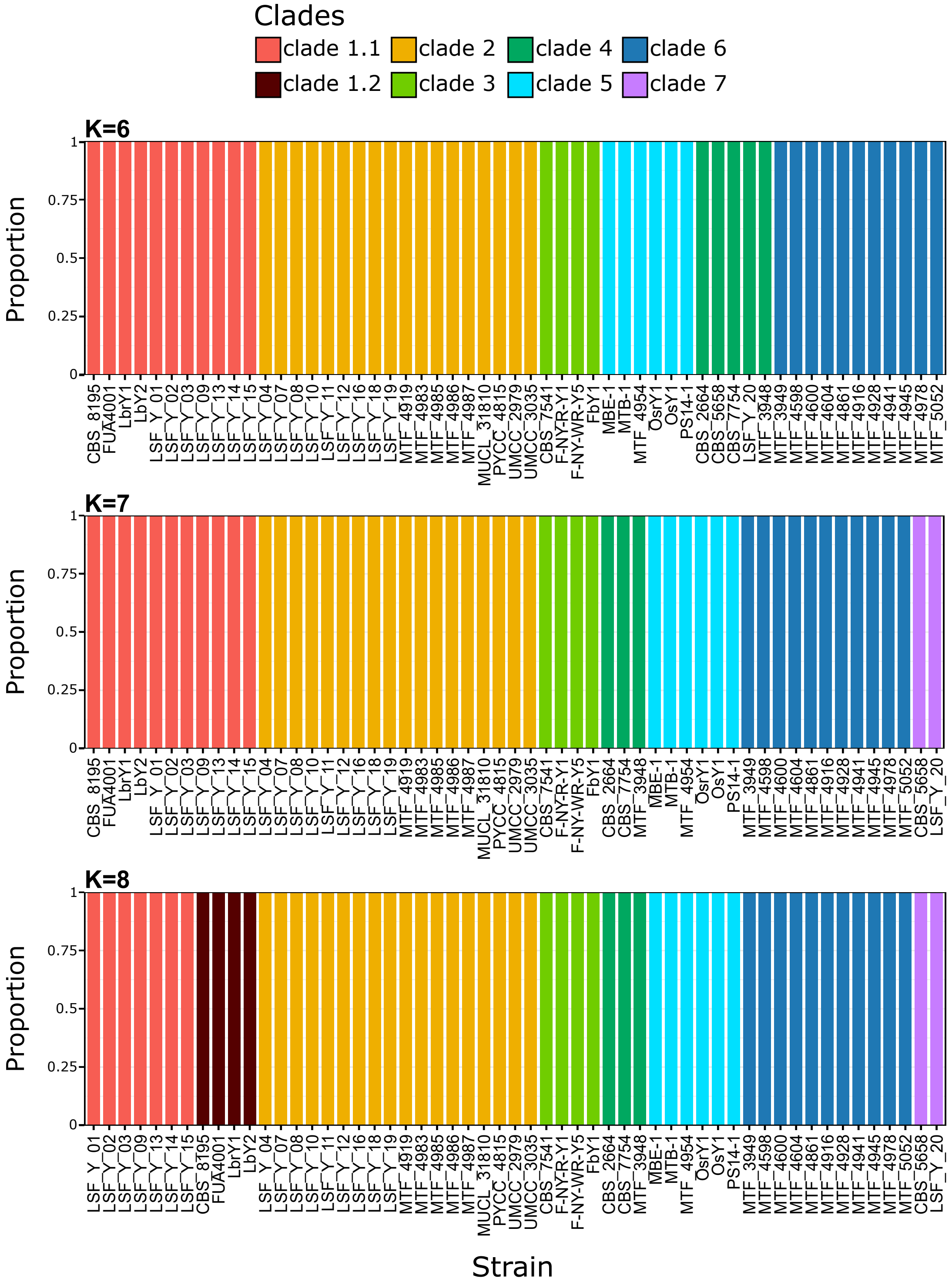


**Fig. S3**: Summary of the discriminant analysis of principal components (DAPC) analysis from 72,561 SNPs and for K = 6 to 8, with clades represented by different colours. Columns represent the estimated allocation to genetic clusters for each strain.

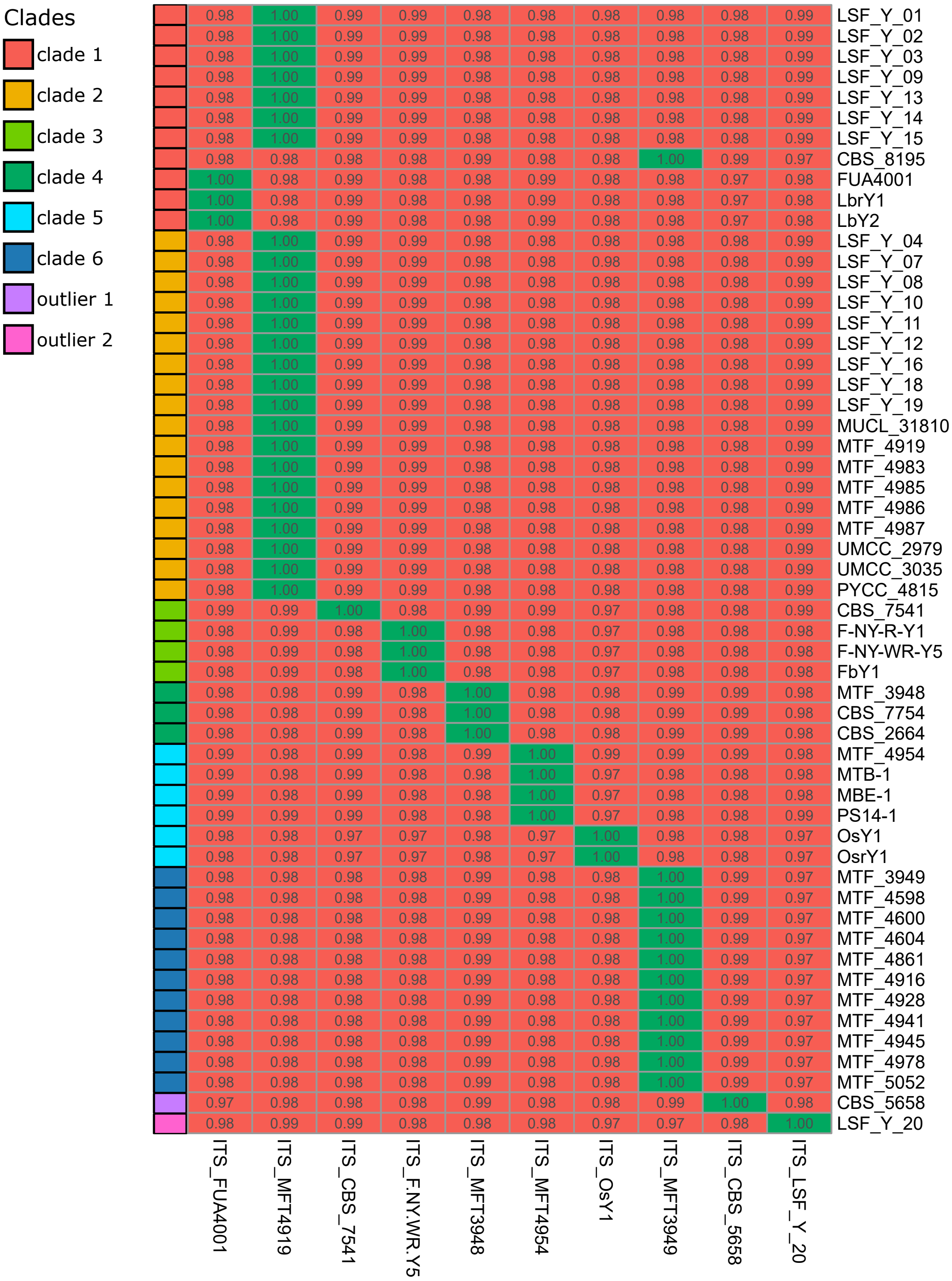


**Fig. S4**: Pairwise identity of the 55 *M. humilis* ITS sequences against the 10 distinct ITS sequences in this species. Identity of 100% is represented in green, lower identity values are represented in red.

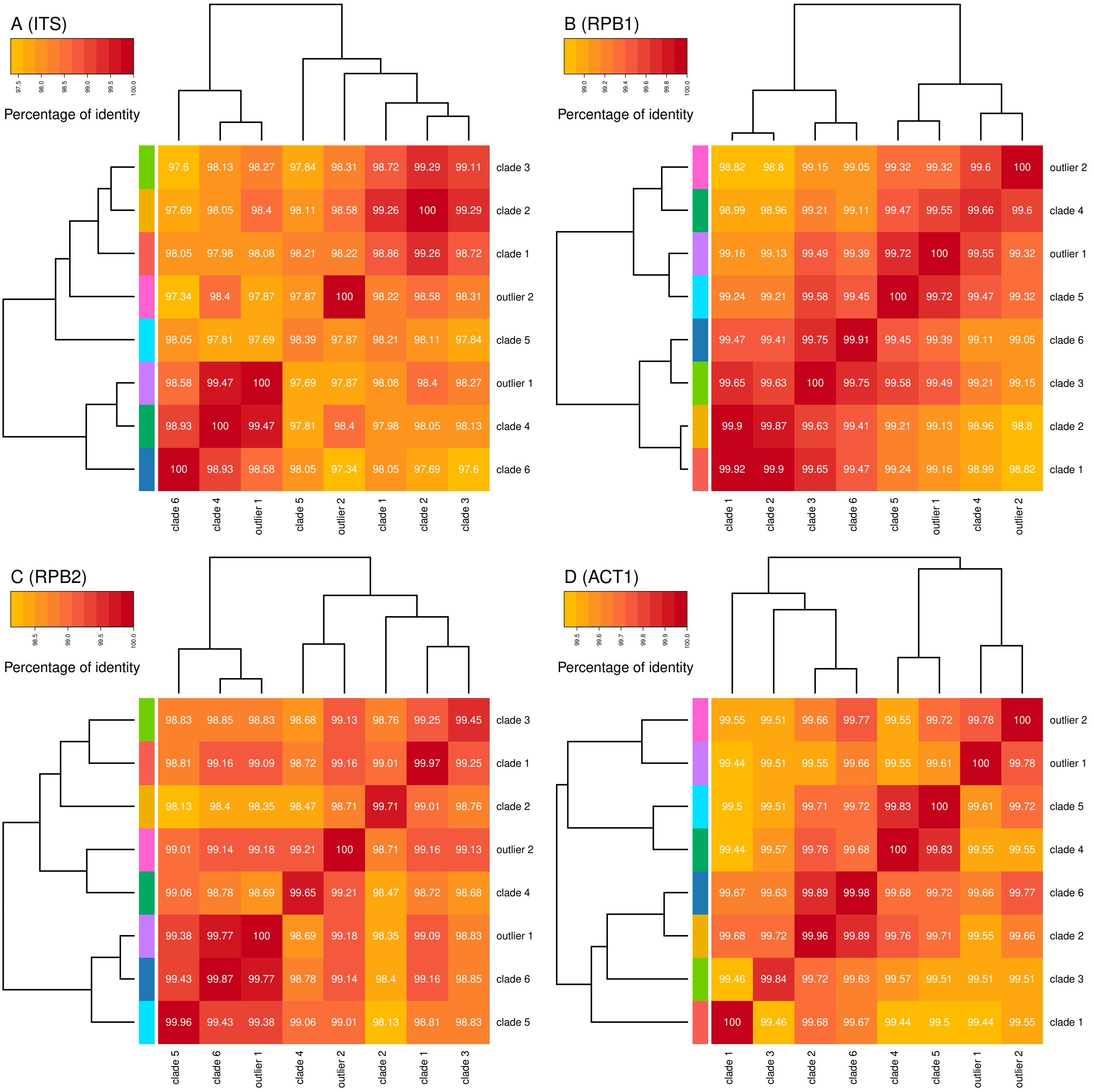


**Fig. S5**: Average pairwise identity of taxonomic markers between *M. humilis* clades. Four markers were considered: ITS (A), RPB1 (B), RPB2 (C) and ACT1 (D). Identity values below 100% on the diagonal indicate polymorphism within clades.

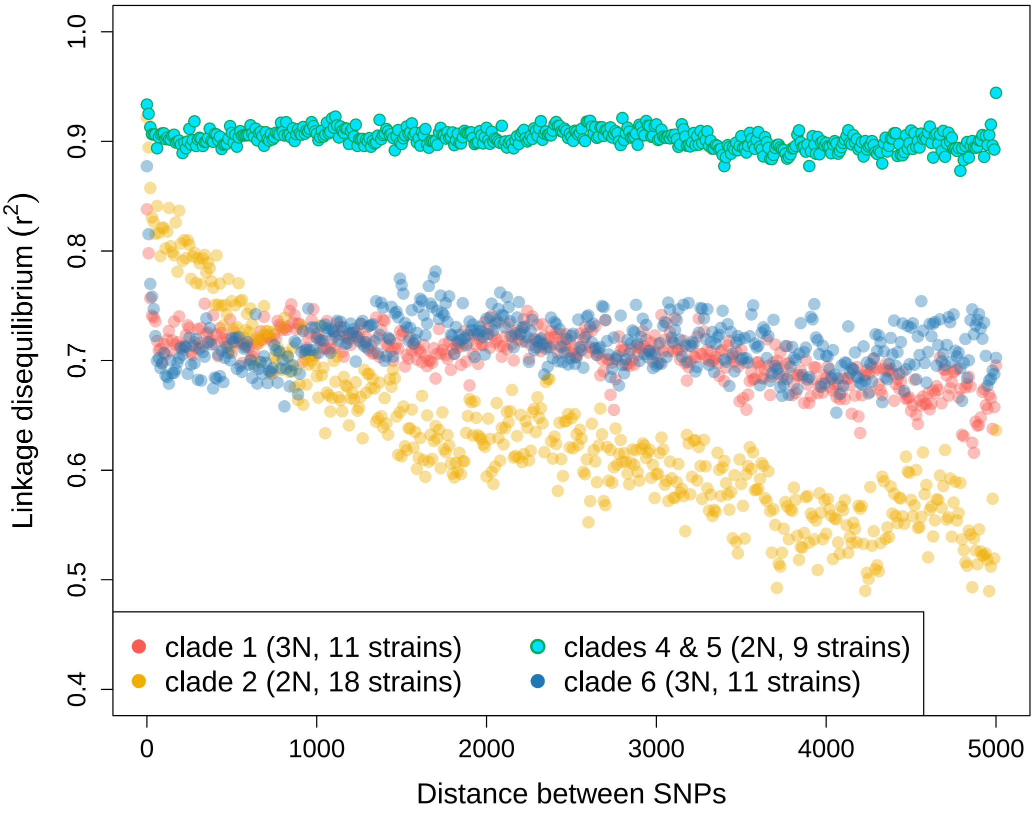


**Fig. S6**: Linkage disequilibrium decay when considering the selection of 72,561 SNPs for clades 1, 2 and 6 separately, and clades 4 and 5 merged.

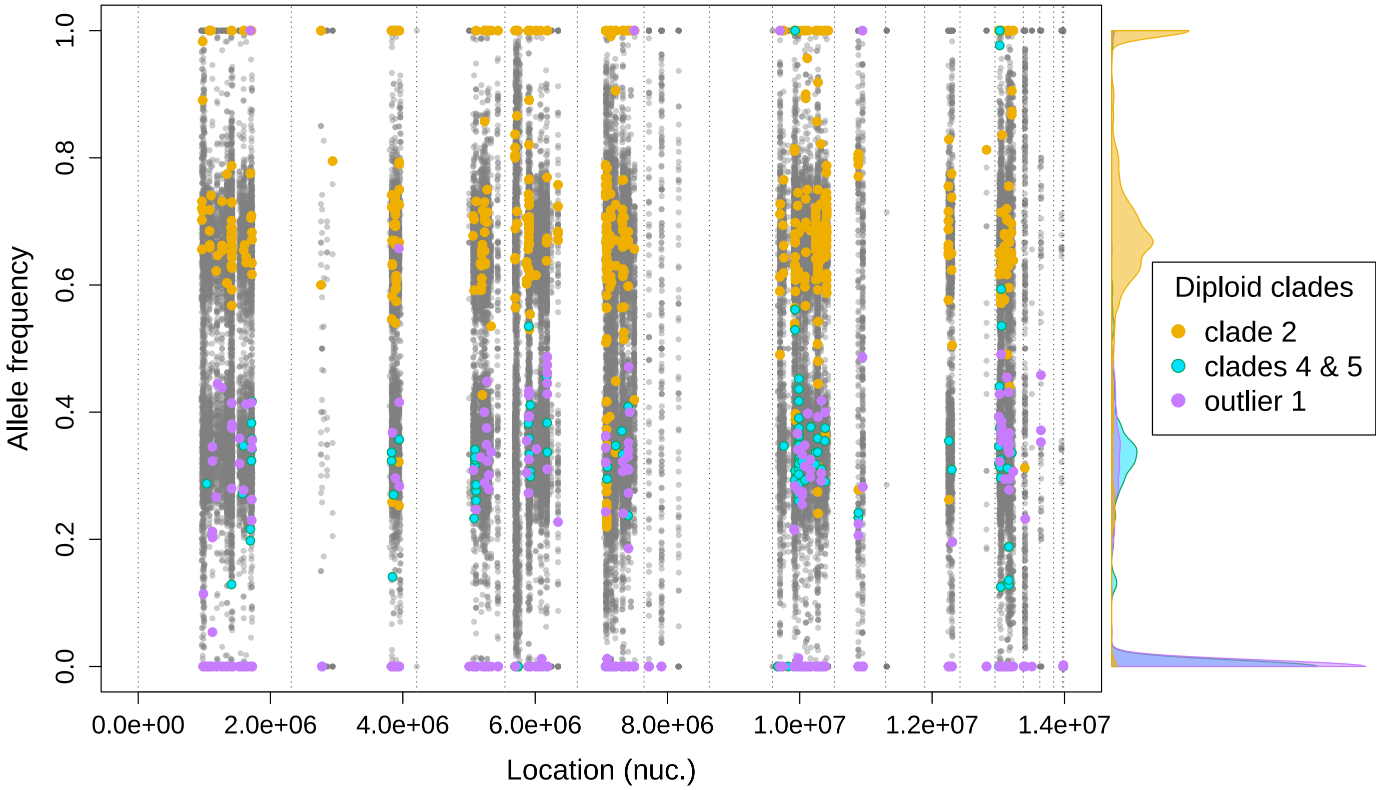


**Fig. S7**: Allele frequencies related to the selection of 72,561 SNPs for the LbrY1 strain (clade 1) reported along the reference genome. Specific homozygous alleles of diploid clades (when considering only diploid clades) are reported as colored dots, clade 2 is in yellow, the merged clades 4 and 5 are in green/blue and outlier 2 is in purple. Allele density as a function of allele frequencies is reported on the right margin, using the same color code for the diploid clades.

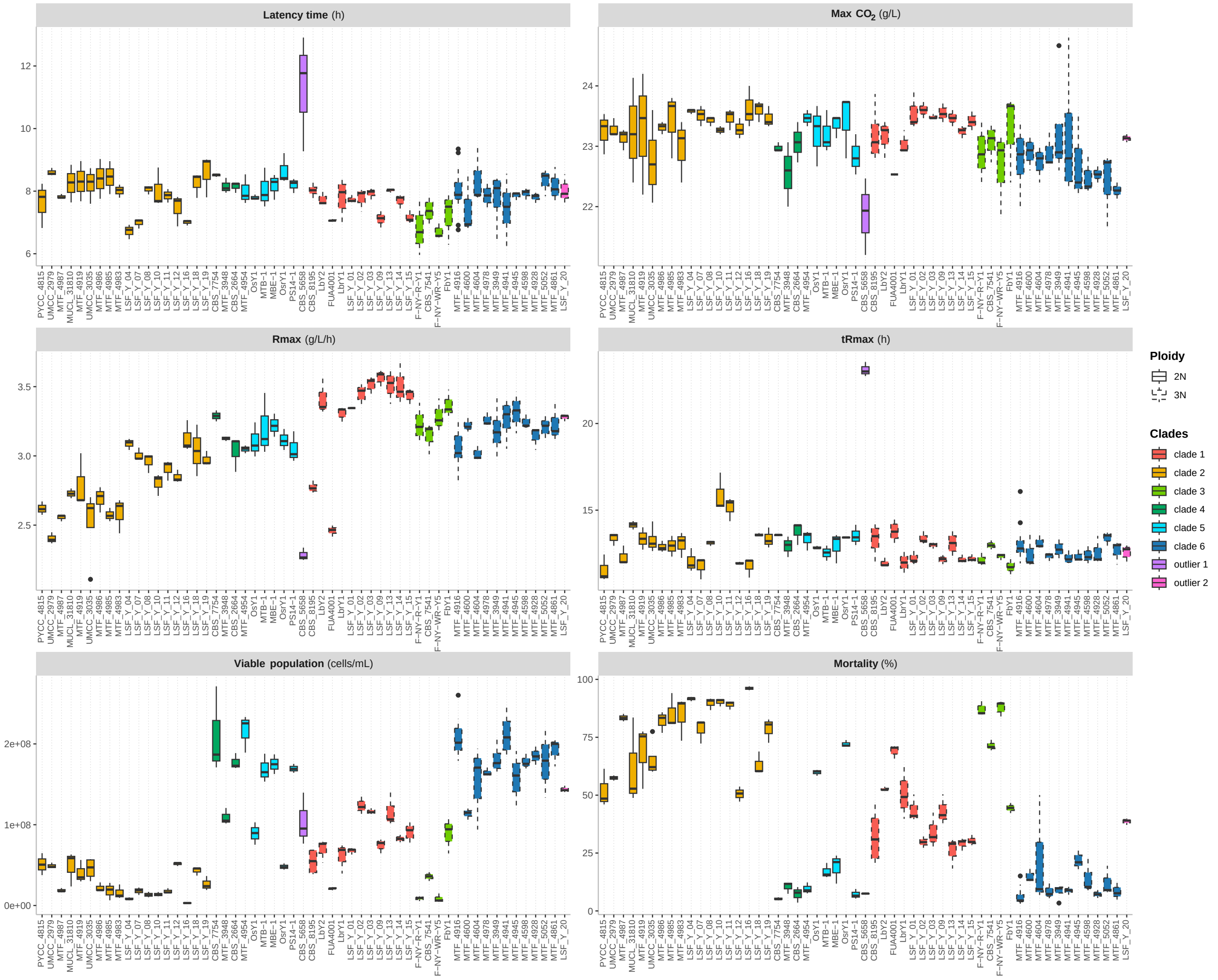


**Fig. S8**: Fermentation performance and fitness in synthetic sourdough medium of 55 strains of textitM. humilis. Fermentation performance was analyzed through four variables: the fermentation latency-phase time (latency time, in h) measured as the time required to reach 1 g/L of $CO_2$ released, the maximum $CO_2$ release (max $CO_2$, in g/L), the maximum $CO_2$ production rate (Rmax, in g/L/h), and the time to reach the maximum $CO_2$ production rate (tRmax, in h). Fitness was estimated at 27h of fermentation by the viable population size (viable population, in cells/mL) and the mortality rate (mortality, in %). Strains were colored according to clades defined by population structure analysis. The outline of the boxplot indicated the ploidy of the strains. At least three independent experiments were carried out for each strain.

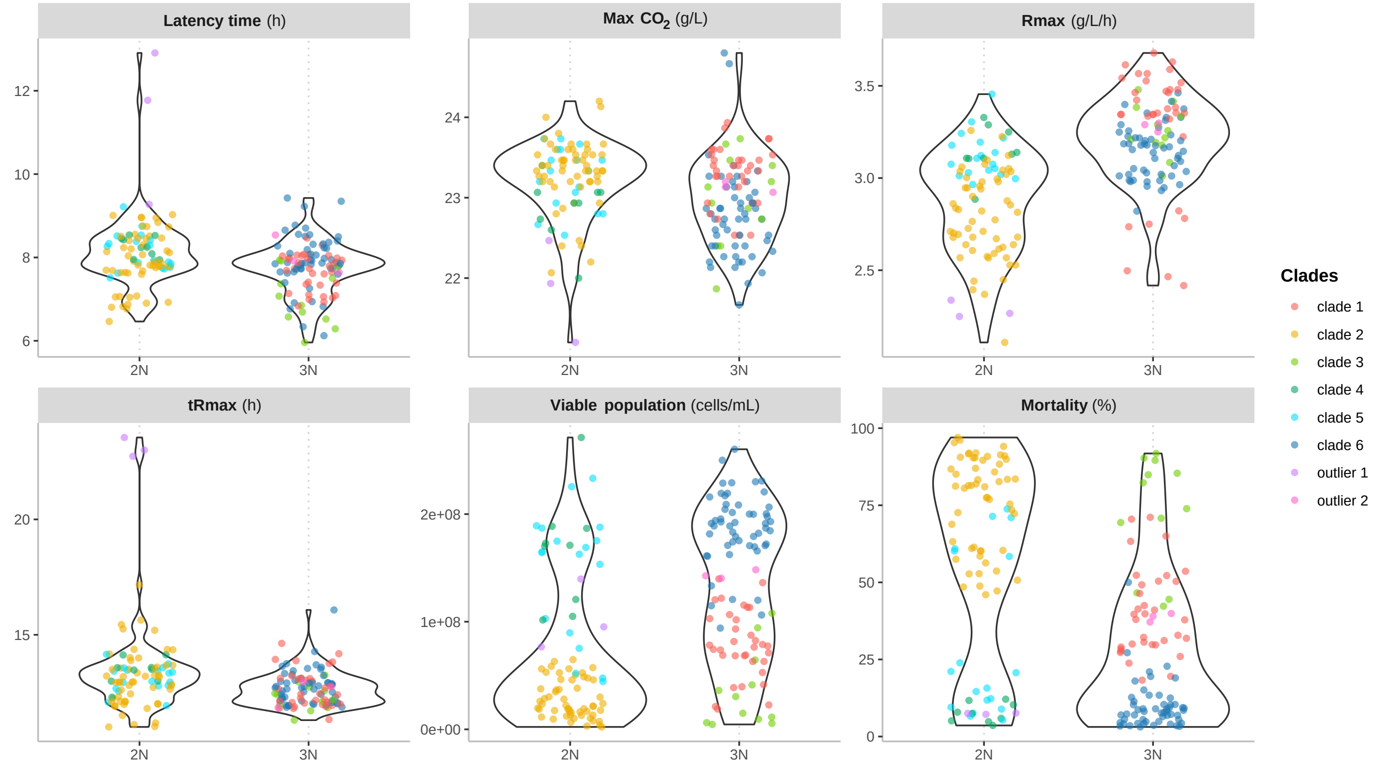


**Fig. S9**: Fermentation performance and fitness in synthetic sourdough medium of 55 strains of *M. humilis* according to ploidy level. For each ploidy, violin diagram was plotted, with observations represented as colored dots according to the clade to which they belong. Each dot represents one fermentation, i.e., one biological repetition. Fermentation performance was analyzed through four variables: the fermentation latency-phase time (latency time, in h) measured as the time required to reach 1 g/L of $CO_2$ released, the maximum $CO_2$ release (max $CO_2$, in g/L), the maximum $CO_2$ production rate (Rmax, in g/L/h), and the time to reach the maximum $CO_2$ production rate (tRmax, in h). Fitness was estimated at 27h of fermentation by the viable population size (viable population, in cells/mL) and the mortality rate (mortality, in %).

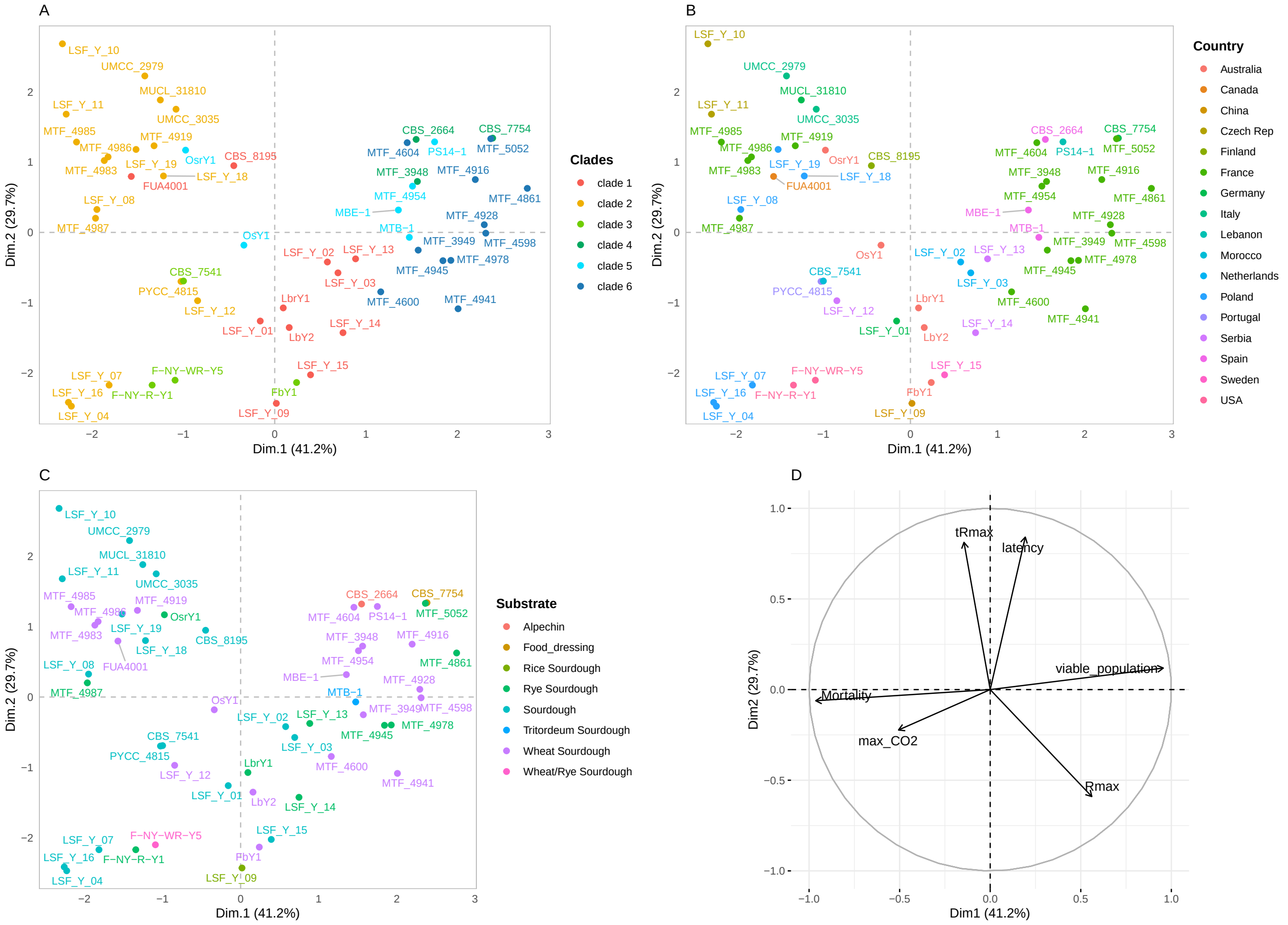


**Fig. S10**: Principal component analysis on fermentation performance and fitness of 53 strains of *M. humilis*. The two principal axes along with their respective percentage of variance explained show the distribution of strains according to (A) clades identified in the ADMIXTURE analysis, (B) country of origin and (C) substrate of isolation. (D) Correlation circle of fermentation performance and fitness variables. Fermentation performance was analyzed through four variables: the fermentation latency-phase time (latency time, in h) measured as the time required to reach 1 g/L of $CO_2$ released, the maximum $CO_2$ release (max $CO_2$, in g/L), the maximum $CO_2$ production rate (Rmax, in g/L/h), and the time to reach the maximum $CO_2$ production rate (tRmax, in h). Fitness was estimated at 27h of fermentation by the viable population size (viable population, in cells/mL) and the mortality rate (mortality, in %). Alpechin is liquid waste from olive oil mills. The “Sourdough” designation indicates that the flour used is unknown.
